# Metabolome-scale genome-wide association studies reveal chemical diversity and genetic control of maize specialized metabolites

**DOI:** 10.1101/450338

**Authors:** Zhou Shaoqun, Karl A. Kremling, Bandillo Nonoy, Richter Annett, Ying K. Zhang, Kevin R. Ahern, Alexander B. Artyukhin, Joshua X. Hui, Frank C. Schroeder, Edward S. Buckler, Jander Georg

**Affiliations:** Boyce Thompson Institute, 533 Tower Road, Ithaca, NY 14853; Plant Biology Section, School of Integrative Plant Science, Cornell University, Ithaca, NY14853; Plant Breeding and Genetics Section, School of Integrative Plant Science, Cornell University, Ithaca, NY14853; Department of Chemistry and Chemical Biology, Cornell University, Ithaca, NY14853; United States Department of Agriculture-Agricultural Research Service, Robert W. Holley Center for Agriculture and Health, Ithaca, New York 14853; Elo Life Systems, 5 Laboratory Drive, Research Triangle Park, NC 27709; Inari Agriculture, Cambridge, MA 02139; Department of Agronomy and Horticulture, University of Nebraska, Lincoln, NE 68583; EAG Laboratories, Columbia, MO 65202

**Author notes:** To whom correspondence should be addressed: Georg Jander, Boyce Thompson Institute, 533 Tower Road, Ithaca, NY 14853. The author responsible for distribution of materials integral to the findings presented in this article in accordance with the policy described in the Instructions for Authors (www.plantcell.org) is: Georg Jander.

## Abstract

**One Sentence Summary:** HPLC-MS metabolite profiling of maize seedlings, in combination with genome-wide association studies, identifies numerous quantitative trait loci that influence the accumulation of foliar metabolites.

**Abstract:** Cultivated maize (*Zea mays*) retains much of the genetic and metabolic diversity of its wild ancestors. Non-targeted HPLC-MS metabolomics using a diverse panel of 264 maize inbred lines identified a bimodal distribution in the prevalence of foliar metabolites. Although 15% of the detected mass features were present in >90% of the inbred lines, the majority were found in <50% of the samples. Whereas leaf bases and tips were differentiated primarily by flavonoid abundance, maize varieties (stiff-stalk, non-stiff-stalk, tropical, sweet corn, and popcorn) were differentiated predominantly by benzoxazinoid metabolites. Genome-wide association studies (GWAS), performed for 3,991 mass features from the leaf tips and leaf bases, showed that 90% have multiple significantly associated loci scattered across the genome. Several quantitative trait locus hotspots in the maize genome regulate the abundance of multiple, often metabolically related mass features. The utility of maize metabolite GWAS was demonstrated by confirming known benzoxazinoid biosynthesis genes, as well as by mapping isomeric variation in the accumulation of phenylpropanoid hydroxycitric acid esters to a single linkage block in a citrate synthase-like gene. Similar to gene expression databases, this metabolomic GWAS dataset constitutes an important public resource for linking maize metabolites with biosynthetic and regulatory genes.

## Introduction

Plants produce wide variety of metabolites that are not directly related to their central energy metabolism and structural integrity. The distribution and diversity of these specialized metabolites are reflective of their essential functions in plant stress responses, especially in their interactions with microbial phytopathogens and insect herbivores. For human societies, plant-derived specialized metabolites have long been valuable sources of flavor, nutrition, and pharmaceutical products. More recently, advances in genetics and molecular biology have led to clarification of the complete biosynthetic pathways of plant specialized metabolites such as glucosinolates (Halkier and Gershenzon, 2006) and benzoxazinoids (Zhou et al., 2018). This knowledge has made it possible to manufacture some plant specialized metabolites at industrial scales, as well as genetically improve crop species for pest and disease resistances.

Productivity of maize (*Zea mays*), the world’s most economically important crop species, with more than 700 million metric tons harvested each year (Ranum et al., 2014), is often limited by pathogens and insect pests (Mueller, 2017). For instance, in parts of Africa, ongoing epidemics of fall armyworm (*Spodoptera frugipeda*) have devastated local maize production, with far-reaching socioeconomic ramifications (Stokstad, 2017). These problems highlight the need for continuous genetic improvement of pest and disease resistance in the commercial maize germplasm to cope with the spatiotemporal fluctuations of biotic stresses. Even after millennia of artificial selection, maize is known for its genetic diversity at the population level (Buckler et al., 2006; Jiao et al., 2017). Similarly, different maize inbred lines possess distinct, tissue-specific profiles of specialized metabolites (Meihls et al., 2013; Wen et al., 2014; Handrick et al., 2016; Wen et al., 2016). Therefore, combining high-throughput metabolite profiling, existing genetic resources, and genotypic data for a metabolome-scale genome-wide association studies (GWAS) will allow large-scale identification of candidate genes and loci involved in maize specialized metabolism, opening up the possibility of harnessing the natural biochemical defenses found in the broader maize germplasm for improved pest and disease resistance. Similar approaches with rice seedling shoots and maize kernels have led to genome-wide identification of metabolic quantitative trait loci (QTL; Wen et al., 2014; Matsuda et al., 2015; Wen et al., 2016).

In this study, we performed liquid chromatography-mass spectrometry (LC-MS) analysis of the tips and bases of the emerging third leaves of maize inbred lines from the Goodman diversity panel (Flint-Garcia et al., 2005). These two tissue types were chosen because: 1) they represented distinct stages of differentiation, and 2) constitutive concentrations of specialized metabolites tend to decrease as plants age (Cambier et al., 2000; Zheng et al., 2015). The Goodman diversity panel contains 282 maize inbred lines belonging to five genetic subpopulations and has been genotyped with over 29 million single nucleotide polymorphism (SNP) markers (Bukowski et al., 2018). More recently, this genetic mapping panel was analyzed by whole transcriptome profiling of eight distinct tissue-environment combinations (Kremling et al., 2018), including the two tissue types used in the current study. Through metabolomic GWAS of maize seedling leaves, we not only provide new insights into the nature of the maize metabolome, but also have developed a public resource that can be used to associate both known metabolites and previously unidentified LC-MS mass features with specific regulatory and biosynthetic loci in the maize genome.

## Results

### Maize seedling leaf specialized metabolome between tissue types and genetic sub-populations

Through reversed-phase HPLC/high-resolution-MS analyses of 50% methanol extracts, which measures a wide range of mid-polarity metabolites, we obtained the specialized metabolome of the leaf bases of 230 inbred lines and the leaf tips of 264 inbred lines. After filtering, more than seven thousand mass features were detected in at least three of the samples (see Materials and Methods; Supplemental Datasets 1 & 2). To parse the natural variation in this dataset, a principal component analysis (PCA) was performed and showed that tissue type explained over 30% of the observed variance (Figure 1A). Two-way analyses of variance (ANOVA) on the same dataset showed that more than 97% of all the mass features analyzed were significantly influenced by tissue type (FDR < 0.05; Figure 1B; Supplemental Dataset 3). In contrast, genetically defined maize population structure did not make a significant contribution to the variance (Figure 1B; Supplemental Dataset 3), and failed to separate in a PCA analysis, even when metabolomics data were analyzed independently within each tissue type (Figure 1C,D). Similarly, PCA within either tissue type showed no systematic bias introduced by the different blocks where each maize inbred line was planted (Supplemental Figure 1).

**Figure 1.**
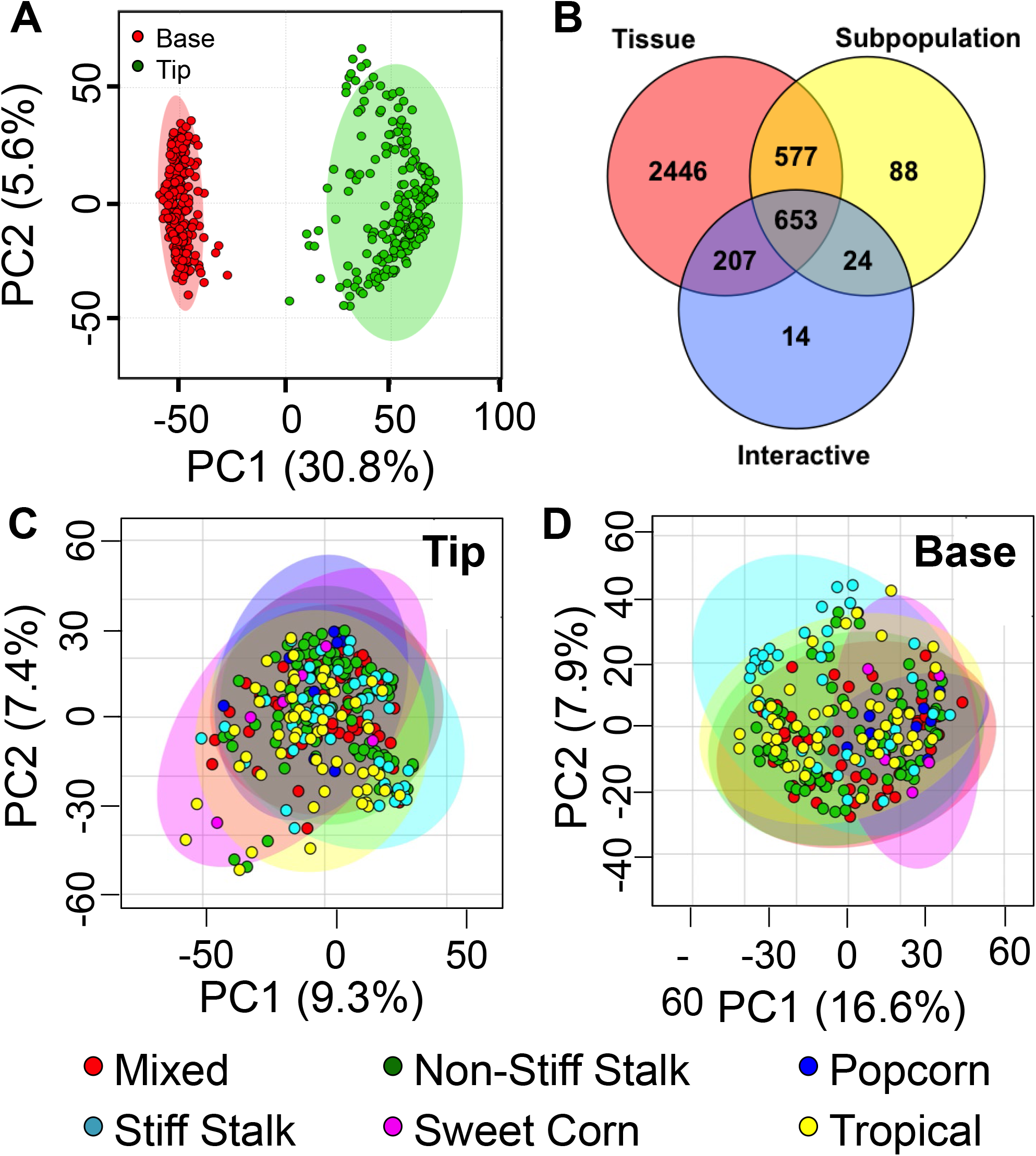
Maize specialized metabolome significantly differentiates leaf tips and bases, but less so among genetic subpopulations. (A) Maize seedling leaf metabolomes can be differentiated between tips and bases with a principal component analysis. (B) Consistently, more mass features are significantly different by tissue type than between subpopulations (two-way ANOVA, FDR < 0.05). Number of mass features differed by tissue type (red), subpopulation (yellow), or their interactive effect (blue) are shown in the colored circles, with overlaps. (C,D) Within either tissue type, genetic subpopulations cannot be differentiated based on their overall metabolomic fingerprint.

### Metabolomic differentiation based on tissue type and genetic subpopulation are driven by different classes of specialized metabolites

In the total ultraviolet (UV) absorption chromatogram, neighboring peaks tended to have similar UV absorbance profiles. Specifically, peaks eluting between 240 and 360 s had UV absorbance profiles resembling phenylpropanoids, peaks eluting between 360 and 460 s had typical benzoxazinoid-like UV absorbance profiles, and those eluting after 460 s were flavonoid-like in their UV absorbance profiles (Supplemental Figure 2). These observations suggested that the range of mass feature retention times could be used to assign them into one of these three major classes.

**Figure 2.**
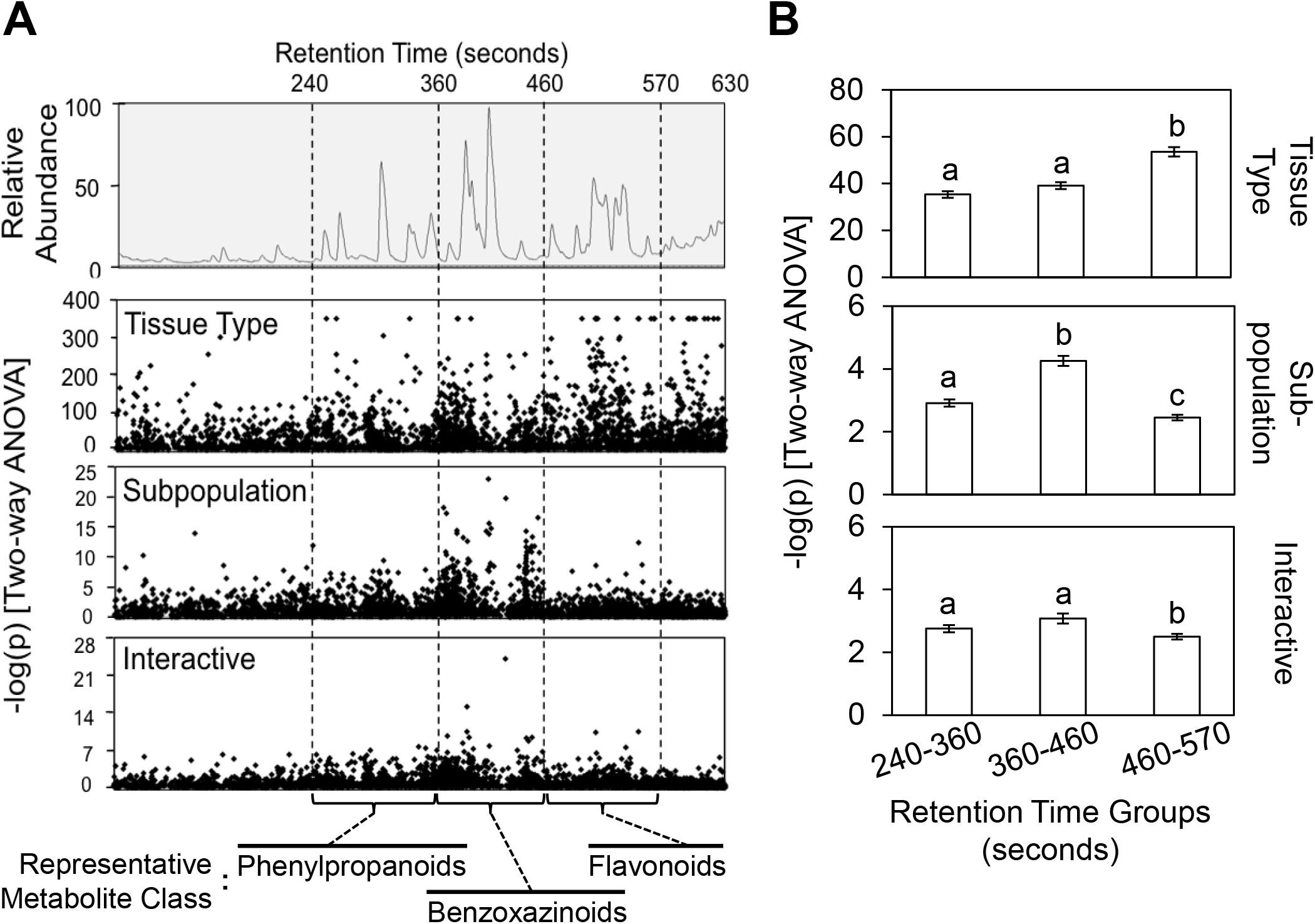
Metabolomic differentiation between tissue types and among subpopulations is driven by different classes of specialized metabolites. (A) Each mass feature is plotted by its retention time (x-axis) and –log(p) by either tissue type, subpopulation, or their interactive effect (y-axes; Two-way ANOVA), and aligned to a sample total ion chromatogram. Representative metabolite class of each retention time range is determined by ultraviolet absorbance profiles of major peaks in the range, as shown in Supplemental Figure 1. (B) Average –log(p) from two-way ANOVA by each variable is compared among three retention time ranges, corresponding to three classes of specialized metabolites. Different letters indicate P < 0.05, ANOVA followed by Tukey’s HSD test. Error bars = standard errors.

We plotted the extent of differentiation for each mass feature based on tissue type, genetic subpopulation, or their interactive effect, as measured by the negative logarithm of p-values from two-way ANOVA, against their retention time (Figure 2A). These plots demonstrated that mass features from distinct ranges of the chromatogram, and hence different classes of specialized metabolites, were responsible for metabolomic differentiation by tissue and subpopulation, respectively. Specifically, mass features that were significantly different between leaf tips and bases tended to fall in the range of flavonoids, whereas those under significant influence from the maize subpopulation or its interaction with tissue type were almost exclusively found among the benzoxazinoids (Figure 2A). These visual patterns were confirmed with statistical comparisons of the extent of differentiation between the retention time groups (Figure 2B). Together, these observations indicate that: 1) flavonoid abundance is significantly different between the maize leaf tip and leaf base, and 2) maize genetic subpopulations differ primarily in their benzoxazinoid content. In support of the first hypothesis, all major mass signals co-eluting with flavonoid-like UV absorption peaks were completely absent in leaf base samples, and only found in the developmentally-advanced leaf tips (Supplemental Figure 3A). Correspondingly, two of the five maize paralogs encoding chalcone synthases, the enzyme catalyzing the first committing step in flavonoid biosynthesis, were differentially expressed amongst the two tissue types (based on B73 reference genome v4; Jiao et al., 2017), and these genes were expressed significantly higher in the leaf tips relative to leaf bases (Supplemental Figure 3B). To test the second hypothesis, we identified mass features representing the most abundant benzoxazinoid compounds in maize seedling leaves, 2,4-dihydroxy-7-methoxy-1,4-benzoxazin-3-one (DIMBOA) and its methylated glucoside derivative, 2-(2-hydroxy-4,7-dimethoxy-1,4-benzoxazin-3-one)-β-D-glucopyranose (HDMBOA-Glc). DIMBOA was significantly depleted in tropical inbred lines, which instead contain significantly more HDMBOA-Glc (P < 0.05, ANOVA), and presumably its highly unstable and undetectable aglucone (Supplemental Figure 4). This pattern is consistent with a previous study of 27 maize inbred lines, which identified a deactivating transposon insertion in *BX12* (formerly *BX10c*), which encodes a DIMBOA-Glc methyltransferase, is more prevalent in tropical than in temperate maize inbred lines (Meihls et al., 2013).

**Figure 3.**
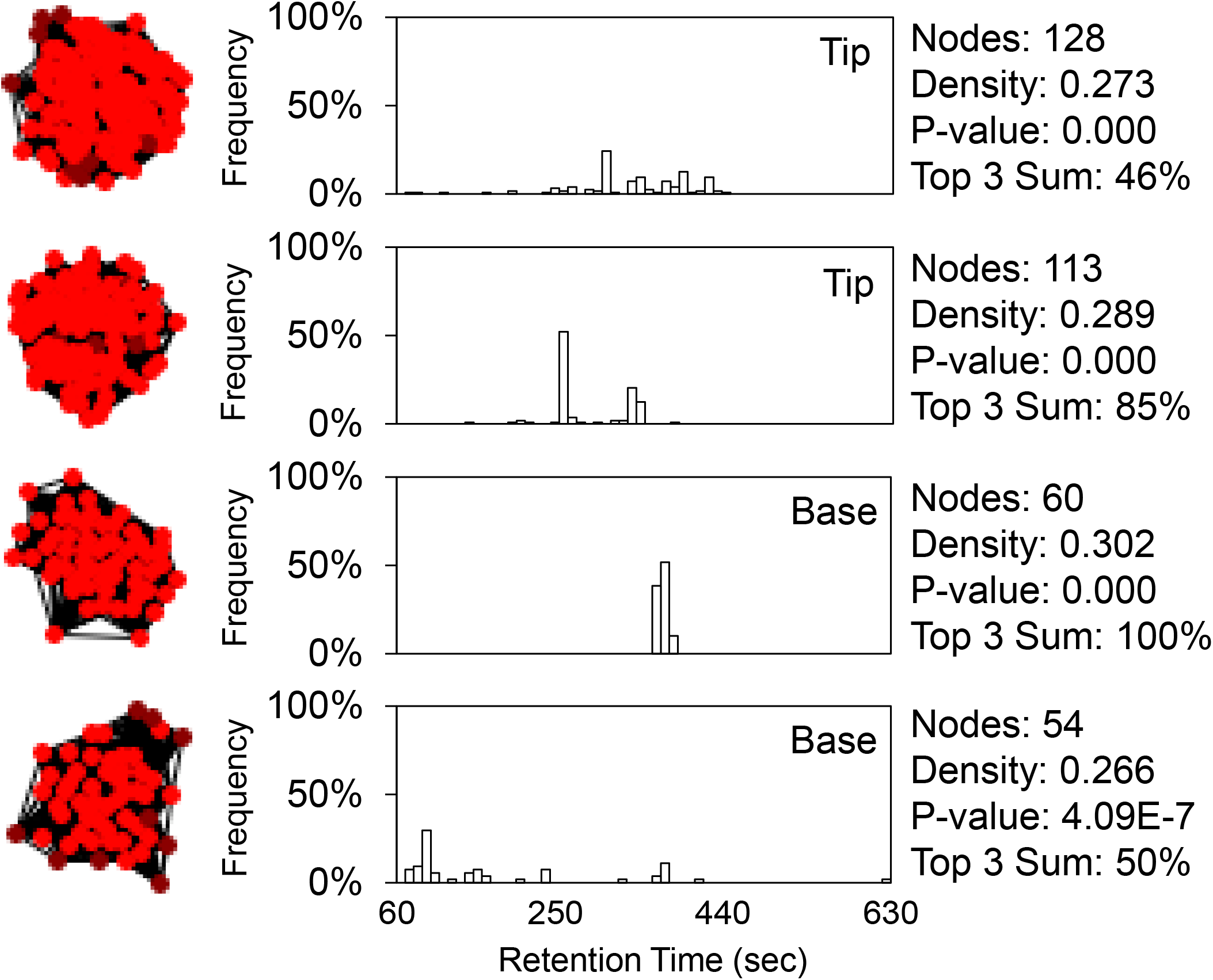
Mass features in the same correlation network tend to have similar retention times. Distributions of retention times of mass features each correlation network are plotted in ten-second increment bins and are aligned to the topographical presentations of the networks. Density and p-value (one-sided Mann-Whitney U test) of each network is calculated by the graph-clustering algorithm ClusterOne. The top 3 sum is the accumulative percentage frequency of the top 3 ten-second bins, which is used to assess the level of clustering in retention time within each network. Only two significant networks with contrasting level of retention time clustering from either tissue type are shown. All other significant network (p < 0.05) are listed in Supplemental Datasets 4 & 5.

### Structurally related metabolites tend to be co-regulated

Since structurally related metabolites usually arise from shared metabolic pathways, we expected these metabolites to be co-regulated in plants, and hence their natural variation should correlate with one another across the population. To test this hypothesis, we constructed mutual rank-based correlation networks with the metabolomic dataset using an exponential decay function (λ = 50), and detected overlapping correlative clusters using the ClusterONE algorithm (Nepusz et al., 2012; Wisecaver et al., 2017). We detected a similar number of significant clusters in leaf tips and bases (p < 0.05, Mann Whitney U test; 15 in leaf tips and 16 in leaf bases). Consistent with the larger number of mass features detected in the leaf tip samples, clusters found in leaf tips were significantly larger than those found in leaf bases (mean = 100 vs. mean = 58; p < 0.005, Student’s *t*-test). The distribution of the retention times of mass features belonging to each correlative network was plotted in 10-second bins, and the extent of retention time clustering of each network was assessed by calculating the cumulative frequency of the top three bins (Figure 3; Supplemental Dataset 4 & 5). In support of our hypothesis, in 24 of the 31 correlative networks detected, at least half of the mass features were located in the top three bins, suggesting that these co-regulated mass features are structurally related. Interestingly, we found that the cumulative frequency of the top three 10-second-bins in correlative networks derived from leaf tip metabolome (57%) was significantly lower than that of leaf base metabolome-derived networks (75%; p < 0.05, Student’s *t-*test).

### Maize metabolome is skewed towards rare metabolites

Our dataset provides an opportunity to examine the diversity of specialized metabolites in maize. In both tissue types, there was a bimodal frequency distribution of the mass feature occurrence rate, as measured by percent of maize genotypes where a mass feature was detected. Whereas 15% of mass features in either tissue type were present in more than 90% of all the genotypes, more than 63% of mass features are found in less than half of the examined genotypes (Figure 4A). If apparently rare mass features are the result of background variation in the MS data set, we would expect them to have a lower signal intensity than mass features resulting from true metabolites. The mean non-zero intensity of each mass feature showed significant positive correlations (R^2^ > 0.96) with its occurrence rate in both tissue types (Figure 4B), suggesting that rare mass features were indeed lower in abundance. However, given the slope of the regression line, a mass feature detected in only 10% of all the genotypes would be on average less than ten-fold lower in intensity than a ubiquitous mass feature. In contrast, mass features of any given occurrence rate show a hundred-fold range in peak intensity (Figure 4B). Therefore, most of the rare mass features are likely to be true maize metabolites that are present in only a subset of tested inbred lines.

**Figure 4.**
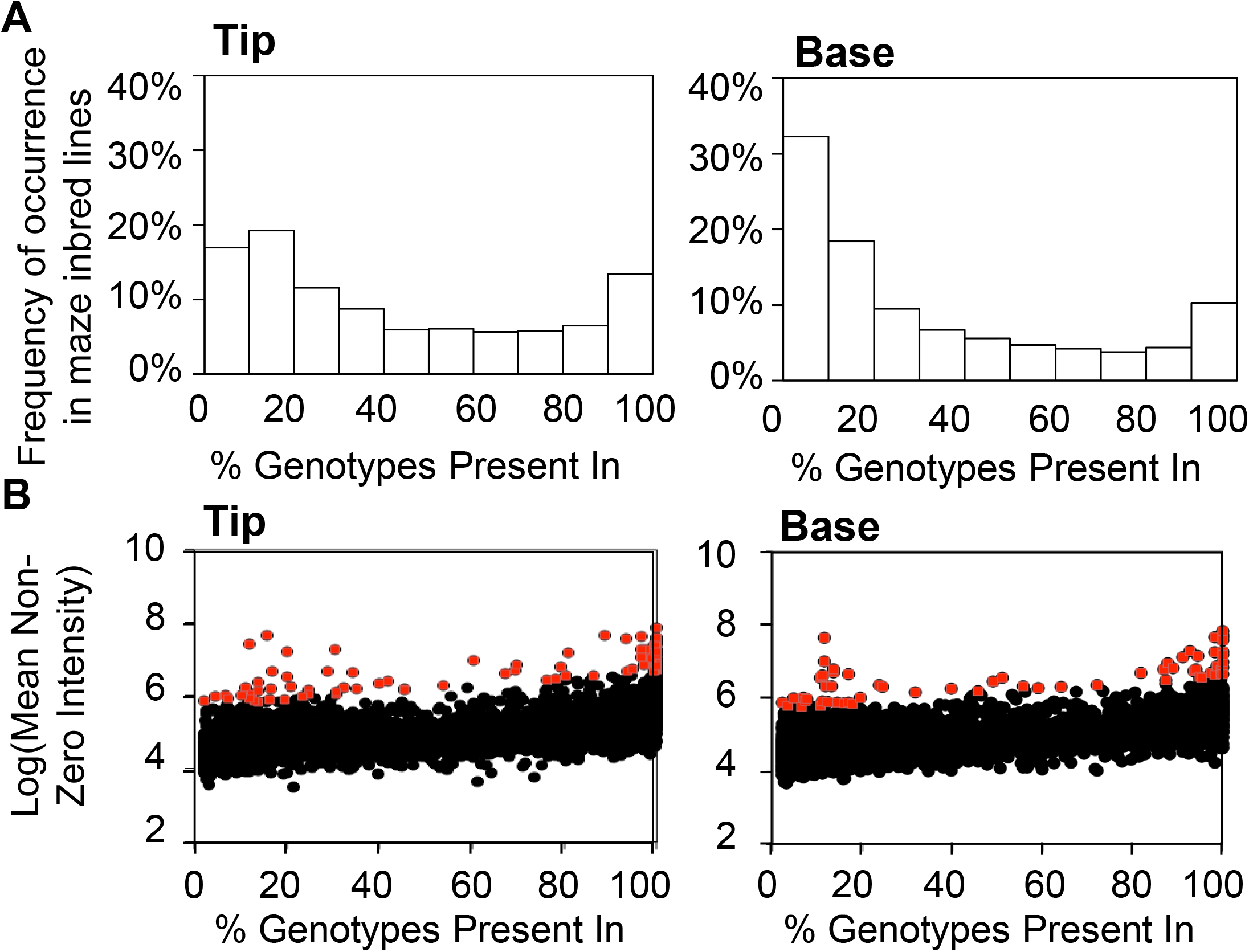
Mass feature occurrence rates are bimodally distributed and are positively correlated with their average non-zero intensity. (A) Distribution of mass feature occurrence rate in either tissue type is plotted in 10% incremental bins. (B) Each mass feature in either tissue type is plotted by its occurrence rate (x-axis) and the log of average non-zero intensity scale (y-axis). Significant positive linear correlations between the two variables are found in both tissue types. Mass features that are above the 99% confidence interval of the overall linear correlation patterns are marker in red.

### Genome-wide association studies with known metabolites reveal both known and novel genetic loci

The existing genotype dataset for the Goodman Diversity Panel (Bukowski et al., 2018) makes it possible to perform GWAS with each mass feature as an independent trait to understand its genetic architecture. Given the large number of traits to be analyzed, a rapid recursive GWAS pipeline was recently developed using an optimized general linear model (Kremling et al., 2018). Two benzoxazinoid compounds, 2-(2,4-dihydroxy-7,8-dimethoxy-1,4-benzoxazin-3-one)-β-d-glucopyranose (DIM2BOA-Glc) and HDMBOA-Glc, were used as positive controls in leaf tips. Biosynthetic genes controlling the synthesis of these two compounds were previously identified through genetic mapping with recombinant inbred populations (Meihls et al., 2013; Handrick et al., 2016). GWAS with both metabolites confirmed the loci containing their respective biosynthetic genes (*Bx10-12* on chromosome 1 for HDMBOA-Glc, and *Bx13* on chromosome 2 for DIM2BOA-Glc), with the most significantly associated SNPs being in linkage disequilibrium (LD) with the respective genes (Figure 5 & 6A). Interestingly, in addition to the SNP markers in LD with the known biosynthetic genes, GWAS also identified SNP markers associated with the metabolites of interest in adjacent LD blocks, suggesting the presence of *cis*-regulatory loci at some distance from the genes of interest (Figure 5 & 6A). Additionally, a locus on chromosome 9 was strongly associated with natural variation in HDMBOA-Glc abundance. At this locus, the most significantly associated SNPs were located within a single 25 kb LD block (Figure 6A). Bi-allelic haplotypes at the mapped loci on chromosome 1 and chromosome 9 were inferred by SNPs within each locus, and inbred lines were assigned to one of two haplotypes using a nearest neighbor cladogram. Haplotype assignment and metabolite quantification, showed an additive effect on HDMBOA-Glc content from *Bx10-12* and the newly identified locus on chromosome 9 (Figure 6B). The single 25 kb LD block on chromosome 9 contained the 3’ region of GRMZM2G108309, a gene model encoding a putative protein phosphatase 2C family protein (Figure 6A). 3’ RNAseq data obtained from seedling leaf tips and bases (Kremling et al., 2018) showed that the expression of GRMZM2G108309 was significantly different between the inbred lines carrying either allele within the LD block (Figure 6C). Furthermore, inbred lines with high GRMZM2G108309 expression accumulated a significantly greater amount of HDMBOA-Glc than those with low expression (Figure 6C). Together, these results suggest that GRMZM2G108309 is a regulator of HDMBOA-Glc content in the tips of maize seedling leaves.

**Figure 5.**
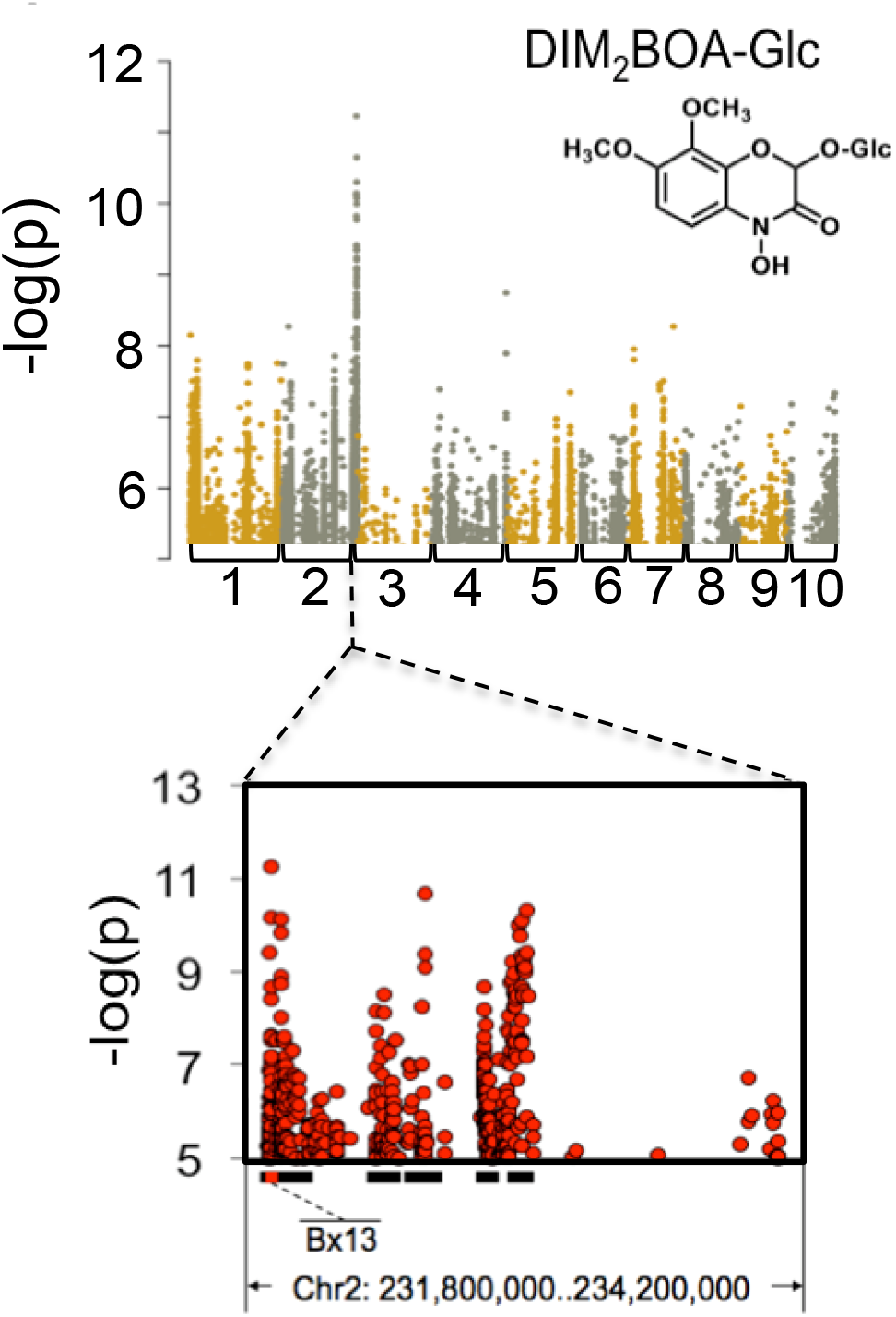
DIM_2_BOA-Glc is significantly associated with genetic markers within and adjacent to its biosynthetic gene. Natural variation in the abundance of DIM_2_BOA-Glc was mapped by GWAS. Each SNP marker is plotted by its physical location in the maize genome (x-axis) and level of association with DIM_2_BOA-Glc abundance (y-axes). SNP markers on adjacent maize chromosomes (labeled on the bottom) are shown in different colors. Only SNP markers with –log(p) > 5 are plotted. Local LD blocks around the most highly associated markers, calculated from the same SNP dataset, are indicated by black bars at the bottom of the plots, and known benzoxazinoid biosynthetic genes are highlighted in red.

**Figure 6.**
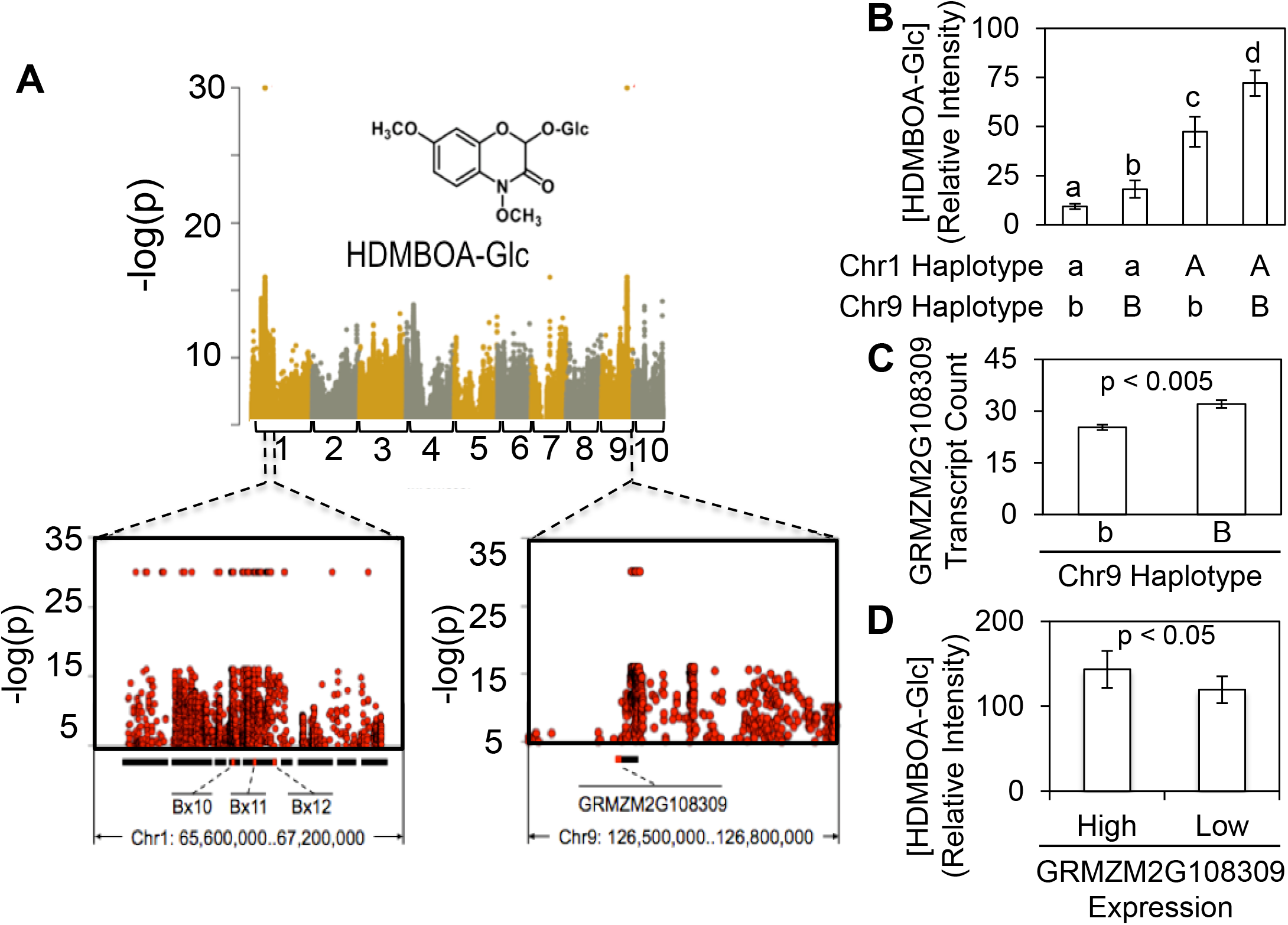
Genome-wide association analysis with HDMBOA-Glc identifies known biosynthetic genes and a previously unknown locus. (A) Natural variation in the abundance of HDMBOA-Glc was mapped by GWAS. Each SNP marker is plotted by its physical location in the maize genome (x-axis) and level of association with HDMBOA-Glc abundance (y-axes). SNP markers perfectly associated with the phenotype (*i.e.* p = 0) were rounded down to 30 on the y-axes for graphical representation. SNP markers on adjacent chromosomes (labeled on the bottom) are shown in different colors. Only SNP markers with –log(p) > 5 are plotted. Local LD blocks around the most highly associated markers, calculated from the same SNP dataset, are indicated by black bars at the bottom of the plots, and known benzoxazinoid biosynthetic genes are highlighted in red. (B) Additive effect on HDMBOA-Glc abundance of the two loci on chromosome 1 and chromosome 9. (C) Effect of haplotypic segregation on the candidate gene expression. (D) Effect of candidate gene expression level on HDMBOA-Glc abundance. Error bars = standard errors.

### Genetic architecture of specialized metabolites is complex and varies independently of heritability and occurrence rate

Based on the successful identification of HDMBOA-Glc and DIM2BOA-Glc biosynthetic loci, we extended our GWAS to the metabolomic scale. Prior to this computation-intensive analysis, the LC-MS dataset was further filtered by the broad sense heritability (*H^2^* ≥ 0.2), and rate of occurrence (detected in ≥ 10% of all genotypes examined). The augmented experimental design allowed estimation of broad sense heritability by calculating the variance of mass features measured in replicated B73 control samples, which were planted in each flat, as the environmental variance. Mass features not detected in B73 but present in other inbred lines were retained, even though their broad sense heritability could not be estimated. Altogether, 1,320 mass features from the leaf bases and 2,554 mass features from the leaf tips remained after these filtering processes (Supplemental Datasets 6-9), and GWAS was performed for each metabolite using a 29 million SNP dataset (Bukowski et al., 2018).

Previous genetic mapping analyses with metabolic traits in rice and maize have often identified small numbers of large-effect genetic loci (Meihls et al., 2013; Chen et al., 2014; Matsuda et al., 2015; Handrick et al., 2016; Wen et al., 2016). To investigate whether these observations represent the rule or the exception in the genetic architecture of metabolic traits, the top 10 most strongly associated SNP markers for each mass feature were collected, and the number of SNPs were counted in 10 kb segments spanning the maize genome. This analysis showed that, in both leaf tips and leaf bases, the ten most significant SNP associations were in an average of 7.4 distinct 10 kb blocks (Figure 7A). If the size of the scanned chromosomal segments was increased to 60 kb or 360 kb (Supplemental Figure 6), the average number of distinct blocks with significant SNP associations decreased to 6.8 and 6.2, respectively, but the overall shape of distribution was not affected. Less than 9% of all mass features analyzed in either tissue type had their top 10 most strongly associated SNP markers located in three or fewer 10 kb blocks. This indicates that metabolic traits, unlike previously assumed, tend to have complex genetic architecture, under control of numerous interacting genetic loci. The most prevalent mass features (occurrence in > 90% of inbred lines) mapped to significantly more loci than mass features that were less prevalent in the population (Figure 7C), suggesting that components of basal metabolism are subject more complex regulation than specialized metabolites. Otherwise, no consistent trend in genetic architecture complexity, as measured by number of mapped loci, was observed with increasing heritability or occurrence rate (Figure 7B,C).

**Figure 7.**
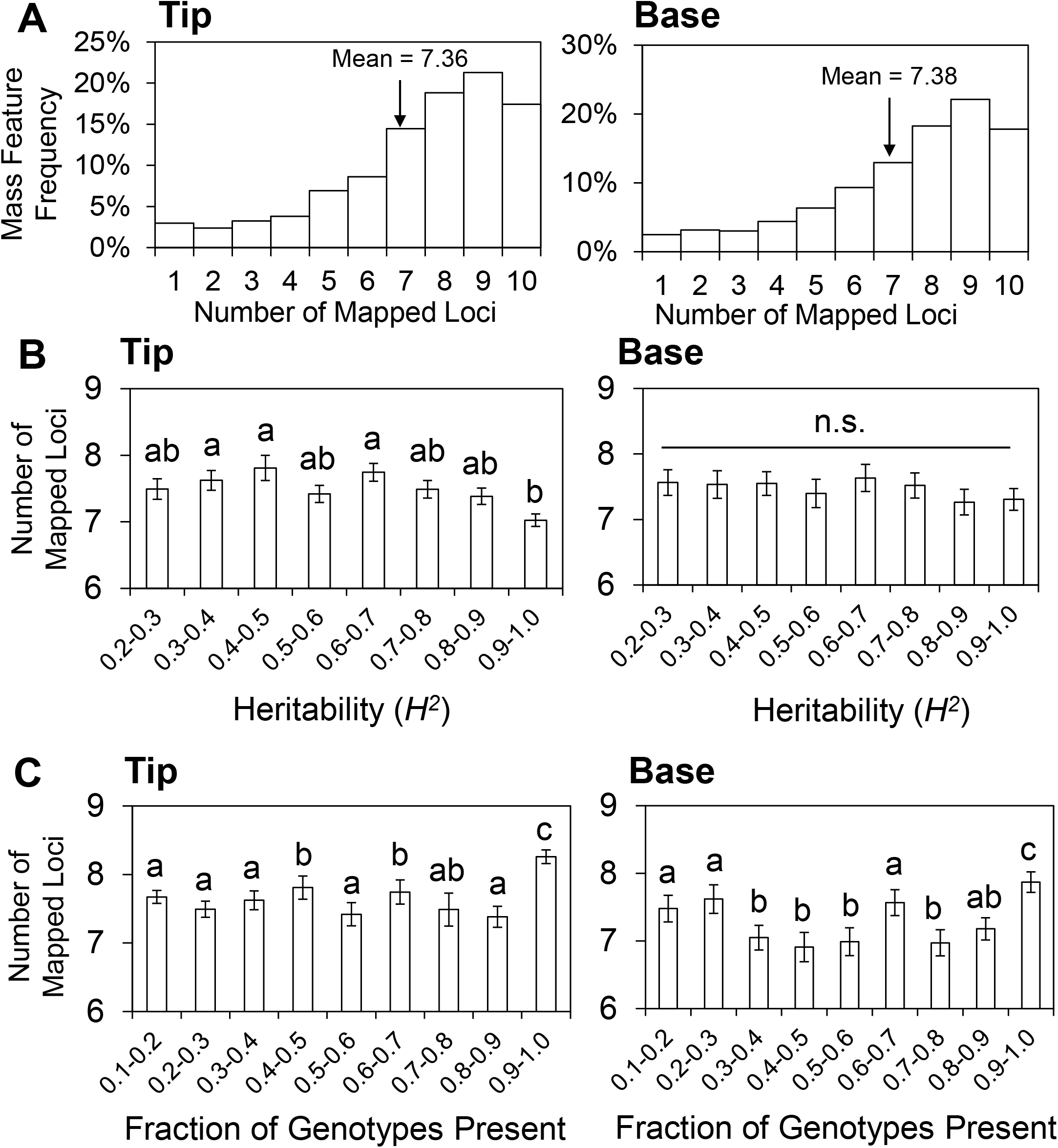
Metabolic traits tend to have complex genetic architecture irrespective of their heritability or occurrence rate. (A) Distribution of mass features in leaf tips and bases is plotted by the number of 10 kb LD blocks that contain one of their top 10 strongest associated SNP markers. Statistical mean of each distribution is given and marked by an arrow. This measurement was then compared across different heritability level (B) and occurrence rate (C) by one-way ANOVA and Tukey HSD. Groups significantly different from each other (p < 0.05) are denoted with different letters on their respective columns. Error bars = standard errors. NS = not significant

### Phenylpropanoid hydroxycitric acid ester isomers found in distinct maize subpopulations are associated with a predicted citrate synthase

A goal of most genetic mapping projects is to identify loci that are significantly associated with specific metabolites of interest, pinpoint candidate genes, and experimentally test for a causal relationship. A metabolome mapping dataset such as the one that we have developed, could be used to rapidly associate LC-MS data with previously unknown biosynthetic or regulatory loci in the maize genome. To test this hypothesis, we genetically mapped the abundance of a relatively under-investigated class of maize metabolites, phenylpropanoid hydroxycitric acid esters (Ozawa et al., 1977; Plenchamp, 2013).

One of the patterns that manifested itself in our analyses of specialized metabolite diversity was that there were clear outliers to the overall positive correlation between the occurrence rate and mean non-zero intensity of mass features (Figure 4B). The majority of these outliers were concentrated in the high occurrence rate range, where the linear correlative relationship was capped by maximal occurrence rate. However, in both leaf tips and bases, a group of high intensity mass features were detected in 20% or fewer of the examined genotypes. Among these outliers, there were three mass features with characteristic phenylpropanoid-like UV absorbance profiles and two common daughter ions with *m/z* = 189.004 and *m/z* = 127.003 under negative electron spray ionization (Supplemental Figure 5A). This suggested that these three peak groups represented conjugates of similar moieties to different phenylpropanoid moieties. Furthermore, the observed mass differences suggested that the phenylpropanoid moieties in these three metabolites differed by hydroxyl and methyl groups, respectively.

In maize inbred lines where these predicted phenylpropanoid metabolites were not detected, at least one other peak was present in each of the three *m/z* channels, all of which had earlier retention times than those that were detected in less than 20% of maize lines. These earlier-eluting peaks also had phenylpropanoid-like UV absorption peaks and had the same daughter ions in MS/MS. The earlier elution times of the peaks with higher occurrence rate suggested that they are structural isomers with higher polarity relative to the three high-abundance peaks that are present in less than 20% of maize lines (Supplemental Figure 5B). To examine how these pairs of structural isomers were distributed, a phylogenetic tree of the Goodman population was estimated using a 66 thousand SNP dataset (derived from those described by Samayoa et al., 2015), and the abundance of the three pairs of structural isomers was plotted across this tree (Figure 8A). In both tissue types, the rare isomers tended to co-occur and were over-represented among the tropical inbred lines. Furthermore, presence of the two groups of isomers was generally mutually exclusive. However, these trends were not perfect, particularly in the case of the isomers with *m/z* = 369.046, both of which were sporadically distributed across the population in the leaf bases without necessarily co-occurring with the other metabolites. The metabolism of these pairs of phenylpropanoid-containing isomers is likely also under developmental regulation, as is demonstrated by the different phylogenetic patterns in leaf tips and leaf bases.

**Figure 8.**
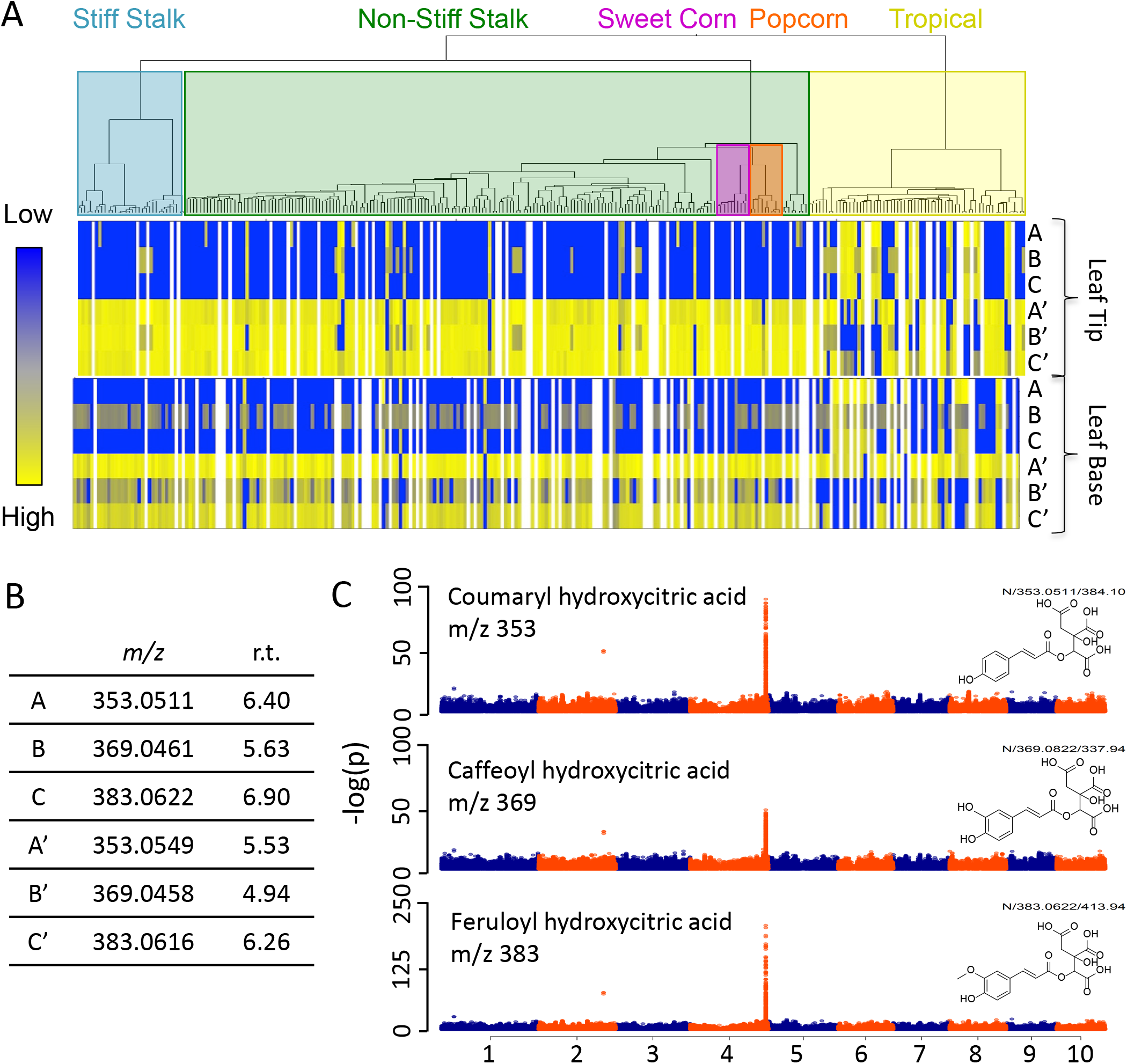
Three pairs of hydroxycitric acid conjugates have complementary distribution and common regulation of abundance in the maize diversity panel. (A) Phylogenetic tree of the 282 maize inbred lines included in the GWAS panel is constructed with the distance matrix calculated from 66 thousand SNP markers. The estimated concentration of three pairs of phenylpropanoid-containing structural isomers are shown in a color scale (blue = low abundance, yellow = high abundance, white = no sample measured) for each maize inbred line. Each monophyletic group is assigned to a genetic subpopulation as defined in Flint-Garcia et al., (2005) by the predominant group assignment for the individuals within that clade. (B) The pairs of mass features shown in panel A have different retention time in minutes (r.t.) were detected in negative ionization mode (*m/z*) and (C) GWAS identified a common locus on chromosome 4 that regulates the abundance of all of the identified mass features from panel A. Only the results from the more polar isomers with structural confirmations are shown.

Two-dimensional nuclear magnetic resonance (NMR) spectra of isolated samples of the more polar isomers that were found primarily in the temperate maize inbred lines showed that they are ester conjugates of coumaric acid, caffeic acid, and ferulic acid, with 2-hydroxycitric acid (Supplemental Dataset 10). In all three cases, these more polar isomers were identified as the 2-O-acylated derivatives. When attempting to isolate the corresponding structural isomers that were primarily found in the tropical inbred lines, the isolated samples rapidly degraded in the NMR solvent, and hence their exact chemical structures could not be fully elucidated. However, their instability suggests that these later-eluting isomers represent the corresponding 3-O-acylated dihydroxycitric acid esters of coumaric acid, caffeic acid, and ferulic acid, given that 3-O-acylated dihydroxycitric acid esters are prone to acid- or base-catalyzed elimination of the 3-O-acyl moiety. Although phenylpropanoid dihydroxycitric acid esters were previously identified as pathogen-induced maize metabolites, their biological function has not been evaluated (Ozawa et al., 1977; Plenchamp, 2013).

GWAS showed that, for all three pairs of phenylpropanoid-hydroxycitric acid ester isomers, the most strongly associated SNP markers were located within a 10 kb LD block on Chromosome 4 (Figure 8C). In the B73 reference genome, this LD block contained a single gene model, GRMZM2G063909, which was annotated as an ortholog of *Arabidopsis thaliana* and *Oryza sativa* citrate synthase family genes (Figure 9). Expression of this gene was not significantly different between maize inbred lines accumulating different structural isomers of phenylpropanoid hydroxycitric acid esters (Kremling, 2018; Supplemental Figure 6), suggesting that structural variation in the encoded enzyme is more likely to be responsible for the observed metabolic differences. To independently verify the genetic association between GRMZM2G063909 and the phenylpropanoid hydroxycitric acid ester isomers, we examined two sets of isogenic lines (NILs) derived from sixth-generation Ki11 x B73 and CML247 x B73 recombinant inbred lines (McMullen et al., 2009) with residual heterozygosity at GRMZM2G063909. Whereas B73 encodes the temperate isomers of the phenylpropanoid hydroxycitric acid esters, Ki11 and CML247 encode the tropical isomers. Both NIL families showed perfect co-segregation between the genotypic markers and the two classes of phenylpropanoid hydroxycitric acid esters (Figure 10A). Furthermore, heterozygote lines showed intermediate phenotypes, producing both isomers, but to lesser abundance than either homozygote. Whereas the two tropical inbred lines also accumulated small amounts of the more polar isomers that are characteristic of temperate inbred lines, B73 tissues did not contain any of the less polar phenylpropanoid hydroxycitric acid esters (Figure 10B,C).

**Figure 9.**
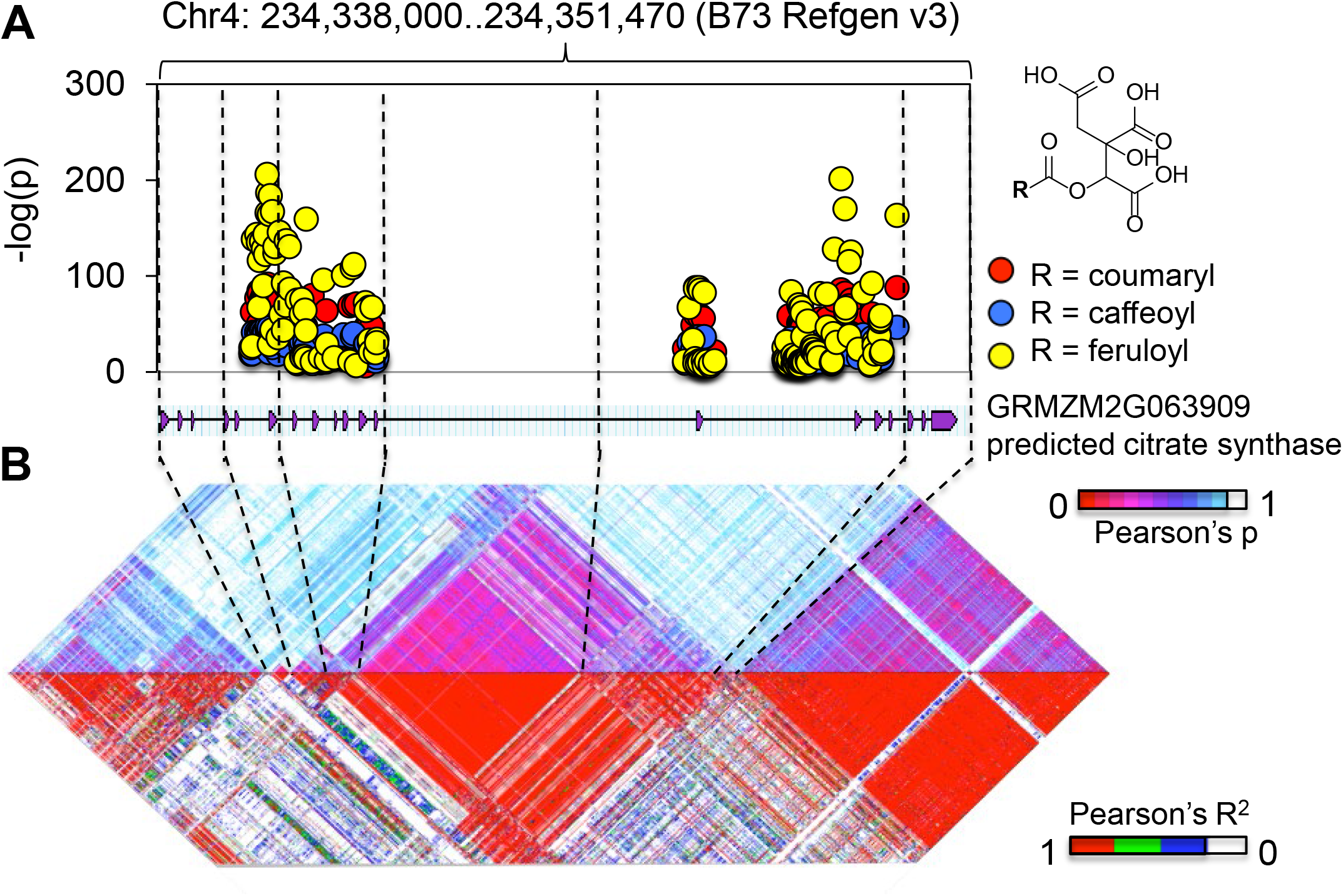
Phenylpropanoid hydroxycitric acid esters are associated with a predicted citrate synthase-like gene. SNP markers most strongly associated with the phenypropanoid hydroxycitric acid esters are plotted by their physical location in the maize genome (x-axis) and level of association with the metabolites (y-axis), and overlaid on the predicted transcripts of GRMZM2G063909 located at the same locus (A). Pairwise correlation coefficients between SNP markers around the candidate gene are calculated to demonstrate that the significantly associated SNP markers are not in linkage disequilibrium with any adjacent gene model (B).

**Figure 10.**
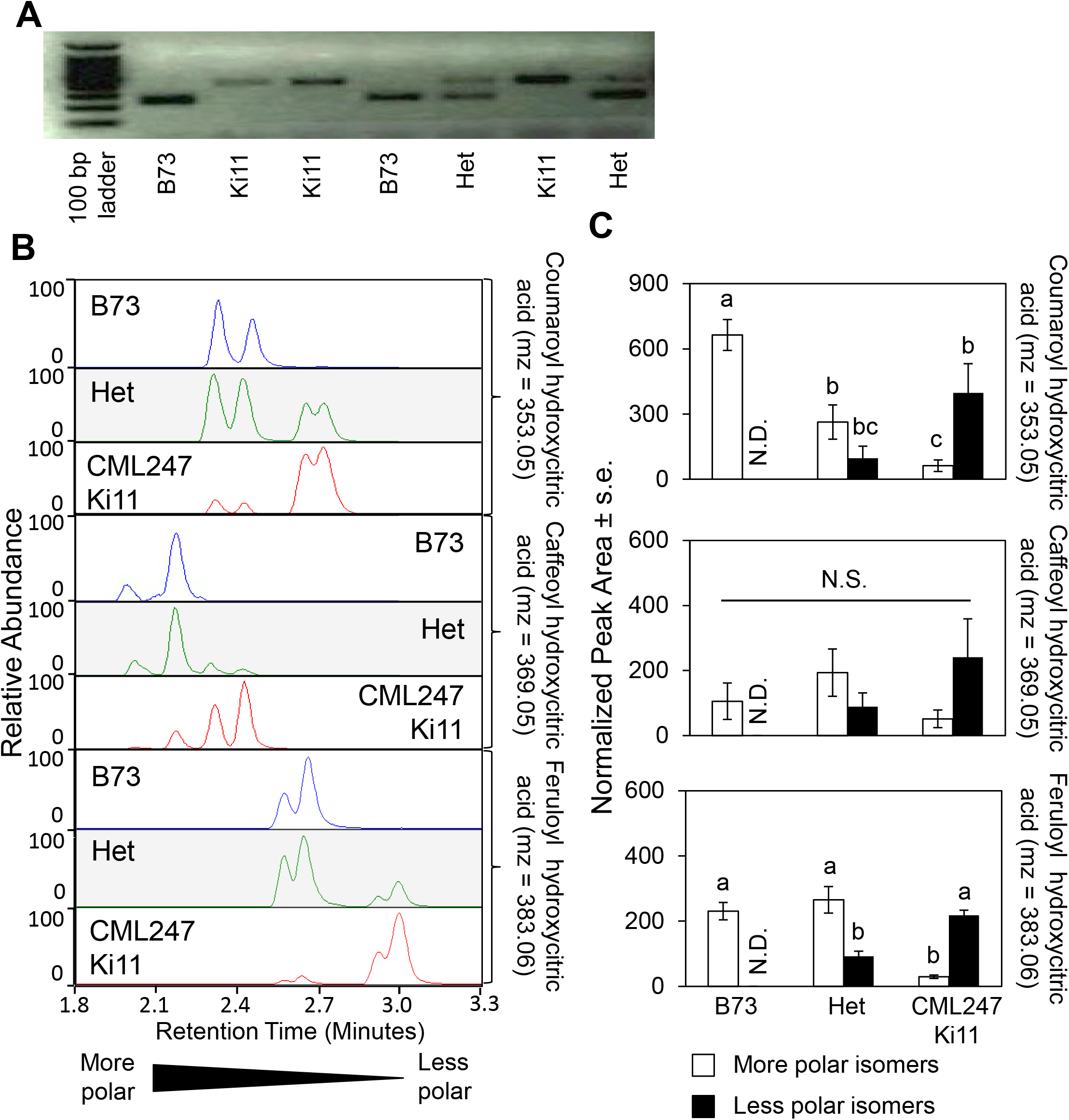
Different isomers of phenylpropanoid hydroxycitric acid esters co-segregate with genetic markers at QTL on Chromosome 4 across near isogenic lines. (A) Representative PCR - based genotyping results and (B) negative ionization mode LC-MS chromatograms of B73 x Ki11 near-isogenic lines. Peak area of all isomers found in near isogenic lines of different genotypes are normalized by total ion concentration of each sample and the total normalized peak area are compared across genotypes and between each other with two-way ANOVA followed by Tukey HSD. Groups of different significance level are indicated by different letters (p < 0.05). N = 14 (B73), 3 (Het), and 4 (CML247/Ki11). N.D. = not detected.

### Structurally related metabolites tend to be co-regulated

In addition to identifying candidate genes significantly associated with individual metabolites of interest, our GWAS results can be used to find genetic loci with disproportionate influence on the overall maize metabolome. When the distributions of the most significantly associated SNP markers for each of the 4,859 mass features were plotted in 10 kbp intervals across the maize genome, this identified several “hotspots” to which a disproportionate number of metabolites were mapped (Figure 11A,C). The locations of these hotspots were consistent when the analysis included either the 10 or the 50 most significantly associated SNP markers for each mass feature, as well as when varying the size of chromosomal blocks used for the plotting the QTL distribution (increasing from 10 to 60 or 360 kbp; Supplemental Figure 7). Thus, the positions of these QTL hotspots are unlikely to be an artifact of the specific data analysis approach.

**Figure 11.**
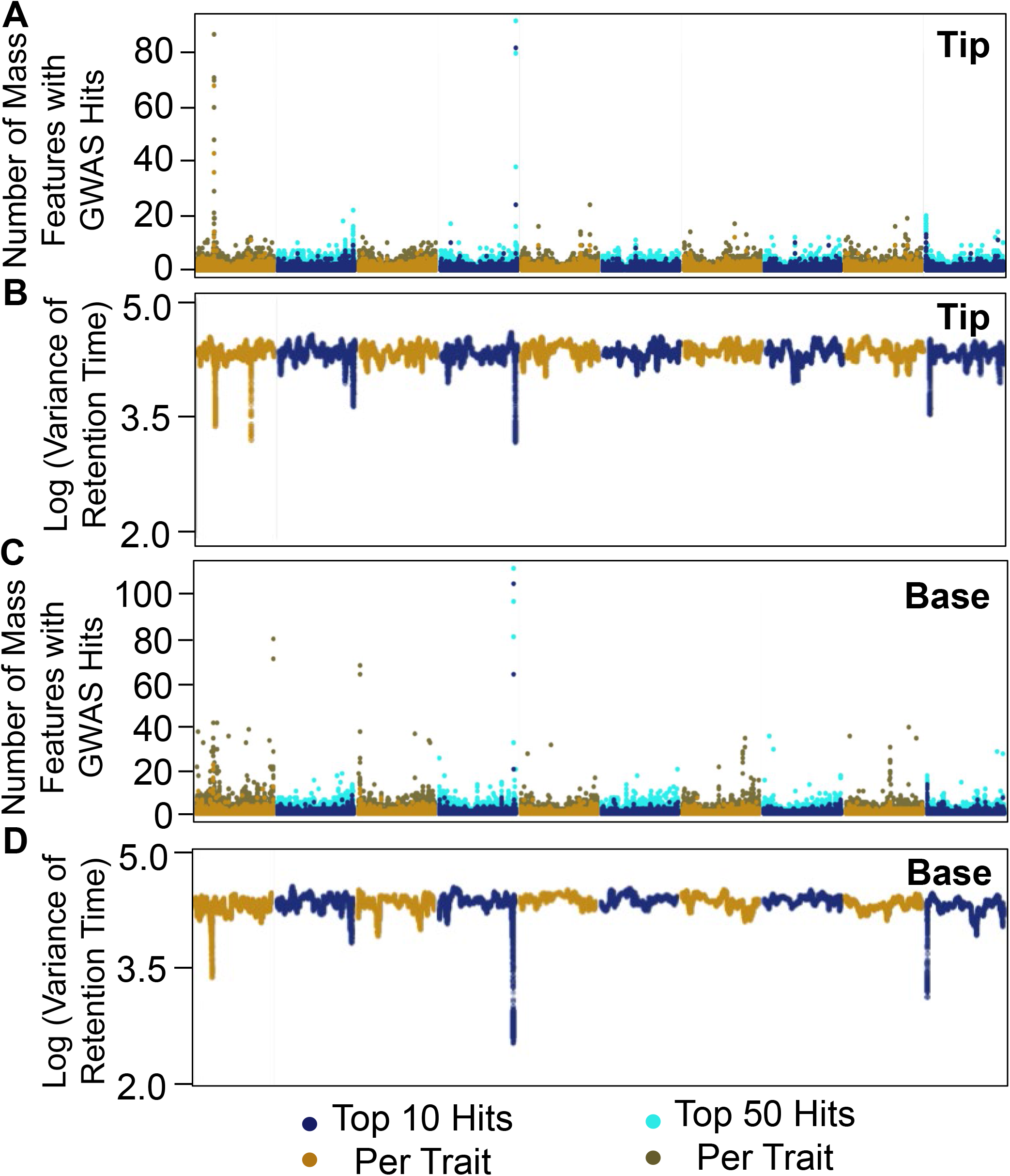
Metabolite GWAS hotspots tend to be associated with mass features that have similar retention times. (A,C) The number of mass features with at least one of their top 10 or top 50 most strongly associated SNP marker located in each 10 kbps block is plotted for either tissue type. Results of neighboring chromosomes are shown in different colors, and results based on different top SNP threshold (10 or 50) are indicated by different color brightness. (B,D) Variance in the retention time of 100 mass features with adjacent GWAS hits in a sliding window across the genome are calculated and mapped based on the physical location of the top SNP hits.

In both leaf bases and tips, three loci on chromosomes 1, 4, and 10, respectively, showed a large number of metabolite GWAS hits. The genomic hotspot on Chromosome 1 represents a 110 kb region containing the *BX11* and *BX12* genes, two paralogous *O*-methyltransferases that catalyze the biosynthesis of HDMBOA-Glc (Meihls et al., 2013). Mass features mapped to this locus include HDMBOA-Glc, DIMBOA, and other benzoxazinoid compounds. Interestingly, many mass features that were not associated with known benzoxazinoid compounds also mapped to this locus, suggesting regulation of other specialized metabolites. Such regulation could be indirect, as benzoxazinoids have been shown to induce other maize defenses (Ahmad et al., 2011; Meihls et al., 2013). The genomic hotspot on chromosome 4, which contained the most GWAS hits in both tissue types, was divided between two 10 kb blocks, one containing the predicted citrate synthase family gene mentioned above (GRMZM2G063909), and the other containing a similarly annotated gene model, GRMZM2G064023. Finally, the less prominent hotspot on chromosome 10, which influenced a dozen mass features in either tissue type, spanned a 30 kb region. In the reference B73 genome, this region contained seven retroelements and a low confidence gene model. The prevalence of transposon genes in this region in the B73 genome suggested that there may be presence/absence variation among the diverse maize inbred lines, and the causative gene may not be present in the B73 reference genome.

In addition to the three hotspots shared between both tissue types, there were also genomic hotspots specific to either tissue type. For instance, 9 mass features found in leaf tips had at least one of their 10 most-associated SNP markers located within a 20 kb region on chromosome 3. This region contained a single gene model, GRMZM2G143723, which is analogous to a rice C2H2 zinc finger protein. These tissue type-specific genomic hotspots were indicative of development-dependent regulation of specialized metabolism in maize seedling leaves.

Among the identified genomic hotspots, two contained confirmed or likely biosynthetic genes. In the case of the chromosome 1 hotspot containing *BX11* and *BX12*, we observed that most of the mass features mapped to this locus represented benzoxazinoid metabolites. This led us to hypothesize that the genomic hotspots contained one or more loci that regulate multiple structurally related metabolites derived from the same biosynthetic pathway. To test this hypothesis, the mass features were ordered by the location of their most strongly associated SNP markers in the maize genome, and the variance in retention time was calculated with a sliding window of one hundred mass features with adjacent QTL in the genome. Since most mass features have their most strongly associated SNP markers at multiple positions in the genome, their retention times were included in the calculations more than once. Across the entire maize genome, there was a stable background level of retention time variance. However, there were clear dips, *i.e.* lower variance in the retention time, below the background level at some loci. When results from this analysis were aligned to the previous plots of mass features per locus, there was co-localization of dips in retention time variance with the genomic QTL hotspots (Figure 11B,D), indicating that abundance of structurally related metabolites tended to be co-regulated by the same genetic loci. This pattern was true for all three of the genomic hotspots shared by both tissue types, but was not necessarily valid all the time. For example, the significant dip in retention time variance on chromosome 1 downstream of the genomic hotspot containing *BX11* and *BX12* did not correspond to any increase in the number of mass features mapped to that locus, whereas mass features mapped to the leaf base-specific hotspot on chromosome 3 did not have similar retention times.

## DISCUSSION

Technological advances in mass spectrometry and accumulating high-density genotypic data are enabling metabolome-scale quantitative genetics studies. Prior studies of this type range from primary metabolites of nutritional interest to known specialized metabolites in both model plants and economically relevant crop species (Chan et al., 2010; Chan et al., 2011; Riedelsheimer et al., 2012; Chen et al., 2014; Wen et al., 2014; Matsuda et al., 2015). However, unlike transcriptomic data, where each transcript can be functionally annotated to at least some extent based on sequence homology and structural features, most mass features from non-targeted metabolomics datasets represent unknown metabolites, and the mass spectrometry data provide incomplete information about their structures. Our metabolome-scale correlation network analyses (Figure 3) and genome-wide association studies (Figure 11; Supplemental Datasets 11 and 12) provide a basis for structural and functional assignments of the many unknown metabolites in maize seedlings. These metabolomic genetic mapping data complement other currently available approaches to metabolite identification, including large scale co-elution tests with known compounds and the construction of molecular networks based on shared tandem mass spectrometry (MS/MS) fragments, which are indicative of structural similarity (Nguyen et al., 2013; Matsuda et al., 2015).

Our datasets allowed us to assess variation in the maize specialized metabolome in two tissue types across a diverse population of inbred lines. The metabolomes of both leaf tips and leaf bases demonstrated bi-modal distributions, with a relatively small core component and a large number of rare mass features (Figure 4A). In comparison to the presence/absence distribution of gene expression in the maize pan-transcriptome (Hirsch et al., 2014), the profiles of our metabolomic data are much more left-skewed, *i.e.* the majority of maize metabolites are present in less than 50% of the inbred lines. This is perhaps reflective of the more commonly non-essential nature of specialized metabolites relative to transcripts, which contain large numbers of housekeeping genes that are involved not only in primary metabolism but also other essential cellular functions. However, the observed distribution differences could also result from the greater sensitivity of RNAseq-based transcriptomics compared to metabolomics, which would allow detection of rare transcripts in a larger number of maize inbred lines.

Our genetic mapping data confirm known associations between benzoxazinoids and their functionally characterized biosynthetic genes (Figures 5 and 6; Meihls et al., 2013; Handrick et al., 2016). The identification of a HDMBOA-Glc regulatory locus on chromosome 9 (Figure 6), which was not found in previous bi-parental mapping studies, highlights the power of broader genetic diversity and denser SNP markers. Additionally, we demonstrated the use of this resource with a group of little-studied metabolites, phenylpropanoid hydroxycitric acid esters (Ozawa et al., 1977; Plenchamp, 2013). Both GWAS and two bi-parental NIL families associated a predicted citrate synthase with isomeric variation in the phenylpropanoid hydroxycitric acid esters that are produced in maize seedlings (Figures 9 and 10). Although, due to their rapid degradation after extraction, we were not able to confirm the chemical structures of the less polar isomers, they plausibly represent the 3-O-acylated isomers, which are prone to decomposition via elimination of the phenylpropionic acid moieties. Structural variation in the identified citrate synthase-like gene is a likely cause of the observed chemical diversity.

In addition to individual genetic locus-metabolic phenotype associations, our study provides a metabolome-scale evaluation of the complex genetic architecture of metabolic traits in maize seedling leaves. Unexpectedly, only a small number of metabolic traits have a simple genetic architecture, as measured by the number of biosynthetic or regulatory loci significantly associated with them. Moreover, we observed no consistent correlative relationship between genetic architecture complexity and heritability or occurrence rate of metabolic traits (Figure 7). We speculate that individual metabolic traits are regulated by different sets of genetic loci in different subsets of the maize population. This observation also could explain the significantly higher number of the mapped loci associated with the most ubiquitous mass features (Figure 7C), which are more likely to be involved in primary metabolism.

Another omic-scale pattern identified from our study are tissue-specific and shared metabolite QTL hotspots (Figure 11). This non-uniform distribution of significant GWAS hits is comparable to results from a published rice metabolite GWAS (Chen et al., 2014). Similar UV absorbance profiles of metabolites in the QTL hotspots indicate that structurally related metabolites tend to be co-regulated by shared genomic loci (Figure 3; Figure 11). The presence of these metabolite QTL hotspots generates hypotheses for the regulation of specialized metabolism both for specific metabolites and at a system scale, and further studies into these loci could lead to elucidation of the underlying physiological mechanisms of these genetic associations.

By demonstrating use of the Goodman diversity panel to map metabolite quantitative traits to the single-gene or near single gene level (Figures 5, 6, and 8), we have generated a rich resource of high-resolution associations between maize metabolic phenotypes and genetic loci. Future researchers who are investigating maize metabolites LC-MS will be able to link their identified mass features with our genetic mapping data to identify potential biosynthetic and regulatory loci. For instance, if our mapping data (Supplemental Datasets 11,12) had been available, the authors who previously reported the discovery of phenylpropanoid hydroxycitric acid esters in maize (Ozawa et al., 1977; Plenchamp, 2013) could have immediately associated their metabolites with GRMZM2G063909, the citrate synthase-like gene that regulates their relative abundance (Figures 8 and 9). Large gene expression data sets generated with DNA microarrays or Illumina-based sequencing (RNAseq) are frequently used for experimental validation and to generate ideas for further research. In a similar manner, our metabolomic association mapping data constitute a community resource that will allow the formulation of testable hypotheses and functional analysis of diverse maize metabolites. Even in the absence of functional validation, the genetic loci and alleles that we have identified will be useful for marker assisted breeding to increase the production of targeted maize metabolites, thereby promoting pathogen resistance or other important agronomic traits.

## MATERIALS AND METHODS

### Plant growth and tissue collection

All maize seeds were originally obtained from the Maize Genetics Cooperation Stock Center (Urbana Champaign, Illinois). To ensure comparability of our metabolomics data with previous published transcriptomics data collected in the same tissue types, the exact same seed stocks were used and the growth conditions was replicated in the same greenhouse space at the same time of the year, early June (Kremling, 2018). Eight seeds of each maize genotype were planted in vermiculite, and the entire diversity panel was fitted into twenty-six 96-cell flats. To control for micro-environmental variation, eight B73 seeds were included in each flat, and all flats were randomized daily. When the third leaf had visibly emerged from the whorl, two centimeters of tissue from the leaf tips and bases were collected. For leaf base tissues, seedlings were cut at the soil line, and unrolled to expose the base. For each maize inbred line, tissues from two seedlings were pooled, weighed, and snap frozen in liquid nitrogen for metabolite extraction.

### Metabolomics analyses and data preprocessing

Frozen seedling leaf tissues were extracted with 200 µL of 50% methanol acidified with 0.1% formic acid, and analyzed on a Sigma Supelco reverse phase C18 column on a Dionex 3000 Ultimate UPLC-diode array detector system coupled to a Thermo Q Exactive mass spectrometer. The two mobile phase solvents were water (Solvent A) and acetonitrile (Solvent B), both acidified with 0.1% formic acid. The mobile phase gradient ran from 95% Solvent A at 0 minutes to 100% Solvent B at 10.5 minutes with curvature of 2 to optimize compound separation while reducing the runtime of each individual analysis to accommodate our large sample size. Each extract was separately analyzed with both positive and negative modes of electron spray ionization. Raw mass spectrometry output files were converted to mzxml formats with the MSConvert tool using an inclusive MS level filter (Chambers et al., 2012). Metabolite quantification was estimated with signal intensity acquired through the XCMS-CAMERA mass scan data processing pipeline (Tautenhahn et al., 2008; Benton et al., 2010; Kuhl et al., 2012). To account for potential rare metabolites occurring in this diverse population, the minimal sample threshold for keeping a mass feature was set at three at the grouping step of the XMCS processing. For initial chemical diversity analyses, LC-MS results from different tissue types were processed together to allow comparison across tissue types. For tissue-type specific statistical analyses and GWAS, only LC-MS results from the same tissue type were aligned to one another and processed as a group to avoid widespread zero values introduced by tissue-specific mass features.

Mass features detected by the XCMS-CAMERA pipeline were filtered based on their retention times (60-630 seconds) and exact masses (*m/z* < 0.5 at first decimal point), and peaks annotated as naturally occurring isotopes were removed. Peaks annotated as MS adducts were retained because we had observed higher rate of false annotation of real metabolites into this category. Mass feature quantification was then corrected by tissue fresh weight and normalized by the total ion concentration of each sample to account for technical variation.

### Chemical diversity analyses

Measurement of each mass feature across the diversity panel was log-transformed for multivariate analyses. Zero values were changed to 1 prior to log-transformation. This dataset was uploaded to the MetaboAnalyst 3.0 online tool platform for principal component analysis and two-way ANOVA (Xia et al., 2015). The mass feature list was further filtered by interquartile range and Pareto scaled before these analyses. In both tissue types, a small number of genotypes had only data available from either positive or negative ionization mode analysis due to failed run under the other mode. These missing data were replaced by zeros to minimize their influence on the overall data structure without losing the usable data. Each maize inbred line was assigned to a genetic subpopulation as defined in Flint-Garcia et al., 2005. All other statistical analyses and data visualization were carried out in R and Microsoft Excel.

### Structural confirmation of phenylpropanoid hydroxycitric acid esters

The three phenylpropanoid hydroxycitric acid esters examined in this study were extracted overnight at 4 °C from bulk snap-frozen B73 seedling leaves with 50% methanol acidified with 0.1% formic acid. Solid debris was removed through centrifugation and the crude extract was concentrated with a Buchi Rotovapor. Target compounds were separated with a water:acetonitrile gradient on a ZORBAX Eclipse XDB C18 column on an Agilent 1100 HPLC system (Agilent, Santa Clara, CA). Purified compounds were dried, weighed, and re-dissolved in pure methanol. NMR spectroscopy analyses were carried out on a Unity INOVA 600 instrument (Varian Medical Systems, Palo Alto, CA) with the following conditions: 256 scans for ^1^H NMR; nt = 16 and ni = 800 for COSY and nt = 32 and ni > 800 for HSQC and HMBC.

### Correlative network analyses

The metabolomic datasets were used to calculate pairwise Pearson correlation matrices, and then mutual rank matrices for the two tissue types separately. Pairwise mutual rank indices were converted to edge weights by an exponential decay functions, with λ = 50 as previously described Wisecaver et al., 2017. For each conversion, edges with weight lower than 0.01 were filtered out. These edge lists were imported into Cytoscape v 3.4.0 (Shannon et al., 2003) and overlapping clusters were detected with the ClusterONE app (Nepusz et al., 2012).

### Genome-wide association study with metabolic traits

The signal intensity of each mass feature across the population was log-transformed. Box-cox transformation was skipped as it distorted the distribution of the rare mass features with lots of zero values. Mass features were filtered based on estimated broad sense heritability and rate of occurrence as described in the Results section, and the remaining 3,991 mass features were analyzed with the fast GWAS pipeline (Kremling et al., 2018). To reduce data storage to a realistic level, only SNPs with – log(p) ≥ 5 for each mass feature were recorded. The top 50 most significantly associated SNP markers of each mass feature from leaf tips (Supplemental Dataset 11) and leaf bases (Supplemental Dataset 12) were extracted for easier reference.

To survey the genetic architectures of metabolic traits, and investigate their relationship with trait heritability and occurrence rate at a metabolomic scale, the top 10 most strongly associated SNP markers for each metabolic trait to three different fixed size LD blocks, namely 10 kb, 60 kb, and 360 kb were mapped. As expected, more metabolic traits have their top GWAS SNP hits located within fewer number of LD blocks as the estimated LD size increases, but the overall shape of distribution was not affected (Supplemental Figure 8). The same LD-block assigning process was used to generate the overview of GWAS hits distribution across the maize genome, by counting the numbers of mass features mapped to each LD block, and plotting them according to the physical location of the LD blocks in the maize genome. Similarly, the locations of metabolite QTL genomic hotspots are consistent across different window sizes of LD (Supplemental Figure 7). Finally, GWAS hits were ordered based on their physical location in the maize genome, and the log variance of mass feature retention time of a hundred adjacent hits was calculated using a sliding window algorithm.

### Local LD estimation, haplotype inference, and inbred lines phylogenetic reconstruction

SNP marker data across the same GWAS diversity panel around the most strongly associated SNP makers for each trait were downloaded from Cyverse Discovery Environment under the following directory (iplant/home/shared/panzea/hapmap3/hmp321), and used to estimate local LD with the pairwise correlation with sliding window algorithm implemented in TASSEL 5.2.40 (Bradbury et al., 2007). Bi-allelic haplotypes at genetic loci associated with HDMBOA-Glc on Chromosome 1 and Chromosome 9 were inferred by SNP data at either locus with a nearest neighbor cladogram also implemented in TASSEL 5.2.40. A smaller SNP dataset (Samayoa et al., 2015) with filtering for maximal missing data (<80%), maximal heterozygosity level (<50%), and minimal minor allele frequency (>30%) was use to estimate the phylogenetic relationship among the maize inbred lines included in this study. Approximately 66 thousand SNP markers were retained after the filtering process and were used to calculate a pairwise distance matrix with TASSEL 5.2.40. This distance matrix was then used to construct a phylogenetic tree using a hierarchical clustering algorithm with the Ward method implemented by the hclust function in R.

## List of Supplementary Materials

**Supplemental Figure 1.** Maize seedling leaf specialized metabolomes are not significantly different among experimental blocks.

**Supplemental Figure 2.** Major peaks from distinct ranges of the chromatogram share characteristic UV absorbance profiles.

**Supplemental Figure 3.** Flavonoids are absent and chalcone synthases expression is low in seedling leaf base tissues.

**Supplemental Figure 4.** Tropical and temperate maize lines accumulate different benzoxazinoid compounds.

**Supplemental Figure 5.** Phenylpropanoid-like mass features co-elute with common daughter ions.

**Supplemental Figure 6.** Expression of the citrate synthase family gene is not significantly different between maize inbred lines accumulating different phenylpropanoid-2-hydroxycitric acid isomers.

**Supplemental Figure 7.** Presence and locations of metabolite QTL hotspots are consistent across different LD window sizes.

**Supplemental Figure 8.** Frequency distributions of genetic architecture complexity of metabolic traits are consistent across different LD window sizes.

**Supplemental Dataset 1.** Mass features detected from positive electron spray ionization mass spectrometry in seedling leaves of diverse maize inbred lines.

**Supplemental Dataset 2.** Mass features detected from negative electron spray ionization mass spectrometry in seedling leaves of diverse maize inbred lines.

**Supplemental Dataset 3.** Two-way analysis of variance results of mass features based on tissue type and genetic subpopulation.

**Supplemental Dataset 4.** Overlapping significant correlative networks of mass features detected in tips and bases of maize seedling leaves.

**Supplemental Dataset 5.** Retention time distribution of mass features in each correlative network.

**Supplemental Dataset 6.** Mass features detected from negative electron spray ionization mass spectrometry in seedling leaves tips of diverse maize inbred lines.

**Supplemental Dataset 7.** Mass features detected from positive electron spray ionization mass spectrometry in seedling leaves tips of diverse maize inbred lines.

**Supplemental Dataset 8.** Mass features detected from negative electron spray ionization mass spectrometry in seedling leaves bases of diverse maize inbred lines.

**Supplemental Dataset 9.** Mass features detected from positive electron spray ionization mass spectrometry in seedling leaves bases of diverse maize inbred lines.

**Supplemental Dataset 10.** 2D-NMR data of maize phenylpropanoid hydroxycitric acid esters.

**Supplemental Dataset 11.** Top 50 most significantly associated SNP markers of mass features detected in maize seedling leaf bases.

**Supplemental Dataset 12.** Top 50 most significantly associated SNP markers of mass features detected in maize seedling leaf tips.

## Acknowledgements

This research was funded by US National Science Foundation awards 1139329 and 1339237 GJ, and US National Science Foundation Graduate Research Fellowship Program award DGE-1650441 to K.A.K. N.B. was funded by the Advanced Research Projects Agency-Energy (ARPA-E), U.S. Department of Energy, under Award Number DEAR0000598. F.C.S. is a Faculty Scholar of the Howard Hughes Medical Institute. This work made use of the Cornell University NMR Facility, which is partially supported by Major Research Instrumentation award CHE-1531632 from the US National Science Foundation. We thank Zack Miller and Lynn Johnson for programming and members of the Jander lab for assistance in plant growth and sample collection.

## Author Contributions

All experiments and analyses were conceived and performed by S.Z. K.A. and J.H. assisted in plant material and LC-MS sample preparations. K.A.K., N.B., and E.S.B. provided advice in experimental design, plant preparation protocols, and performed bioinformatic analysis. A.R. performed experiments with near-isogenic lines. Y.K.Z., A.B.A., and F.C.S. performed 2D-NMR spectroscopic analyses. G.J. provided advice in all experiments and analyses. S.Z. and G.J. prepared the manuscript.

---

**Supplemental Figure 1.**
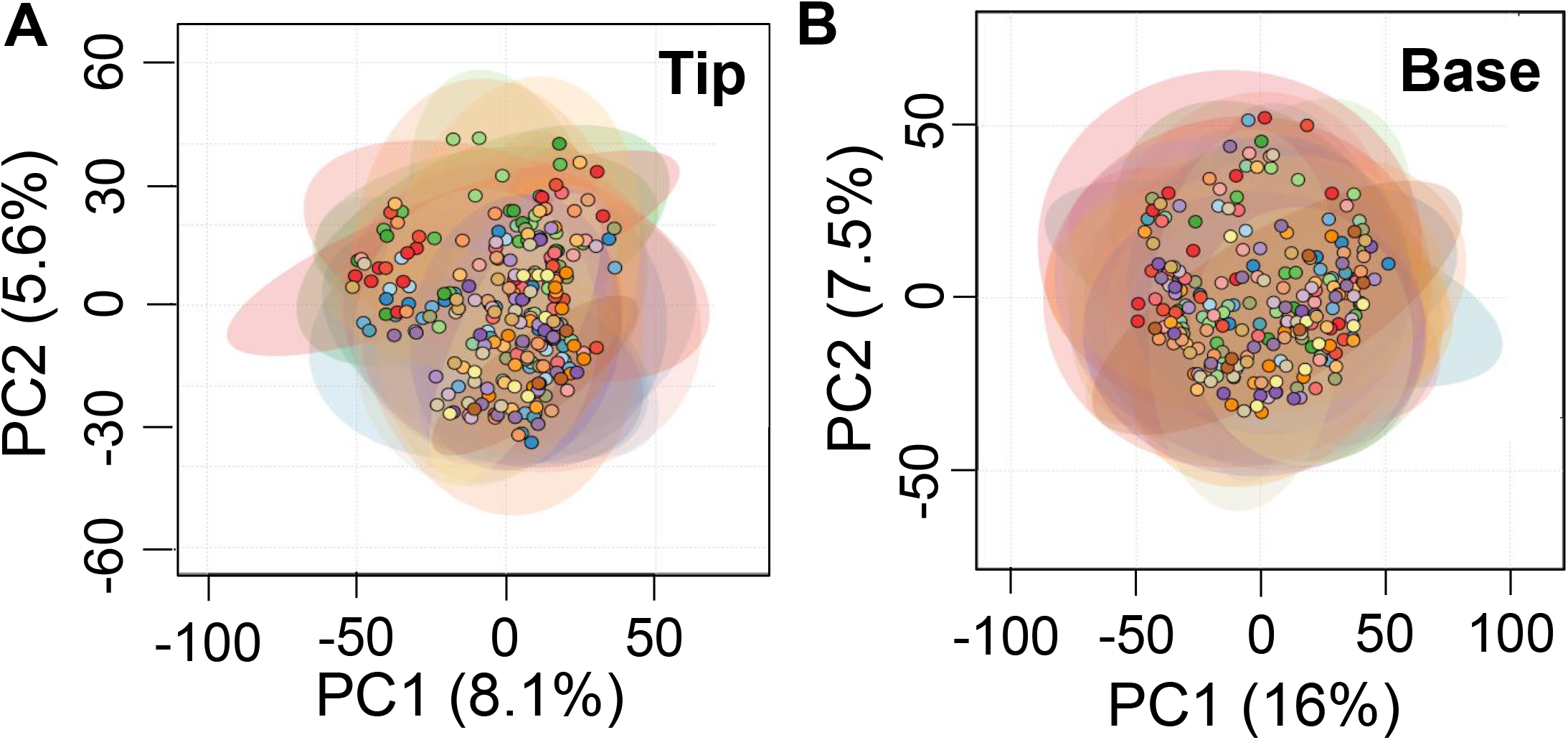
Maize seedling leaf specialized metabolomes are not significantly different among experimental blocks. Principal component analyses of specialized metabolome data collected from (A) tips and (B) bases of the emerging third leaves of diverse maize inbred lines. Each maize line is represented by a dot, and the random experimental block where it is assigned is denoted by different colors. 95% confidence range of each group is shown by the correspondingly colored circles.

**Supplemental Figure 2.**
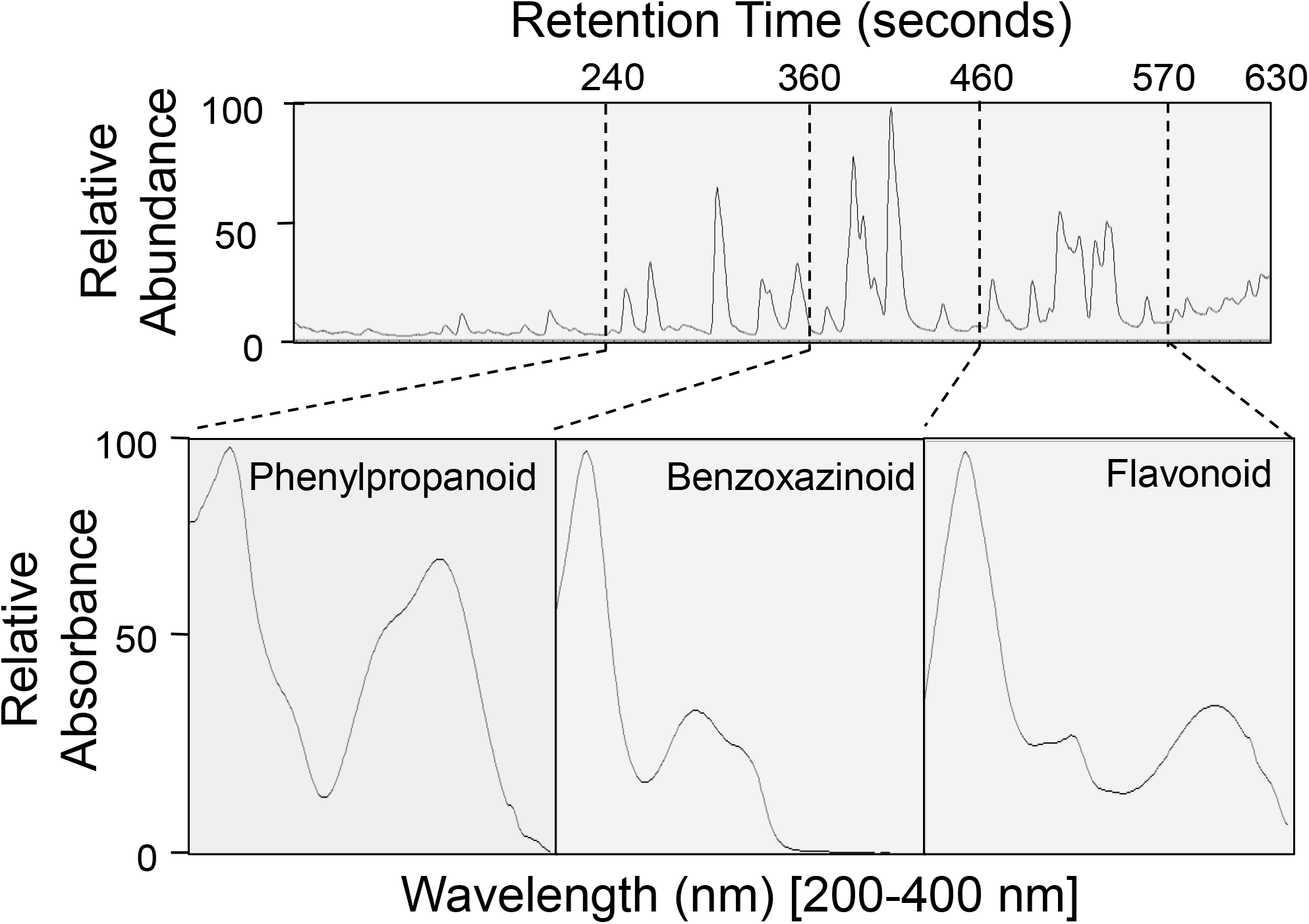
Major peaks from distinct ranges of the chromatogram share characteristic UV absorbance profiles. The UV absorbance profile of each peak was constructed with a photodiode array detector. The boundary of each retention time range was determined by the last peak showing the same characteristic UV absorbance profile. There was no detectable overlap between neighboring retention time ranges.

**Supplemental Figure 3.**
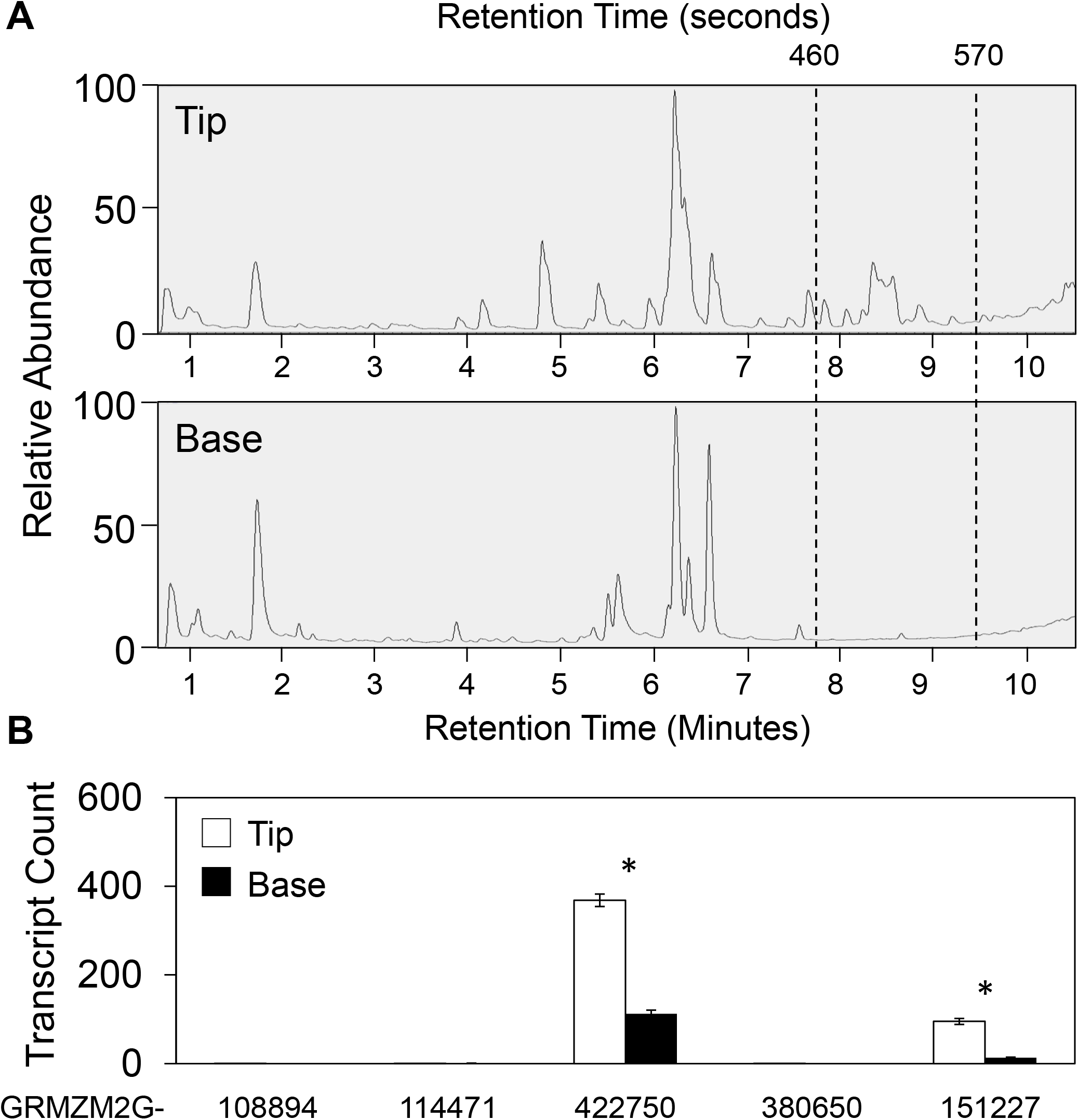
Flavonoids are absent and chalcone synthases expression is low in seedling leaf base tissues. (A) Sample UV absorbance chromatograms of the seedling leaf tip and base of the same genotype are shown to demonstrate the lack of peaks in the flavonoid time range (460-570 seconds). (B) Average expression of five chalcone synthase-encoding gene models in the B73 reference genome v4 (Annotation 5b+) across the Goodman diversity panel are compared between these two tissue types with Student’s *t-*tests (*FDR < 0.05). Error bars = standard errors. Expression data were obtained from Kremling et al., 2018.

**Supplemental Figure 4.**
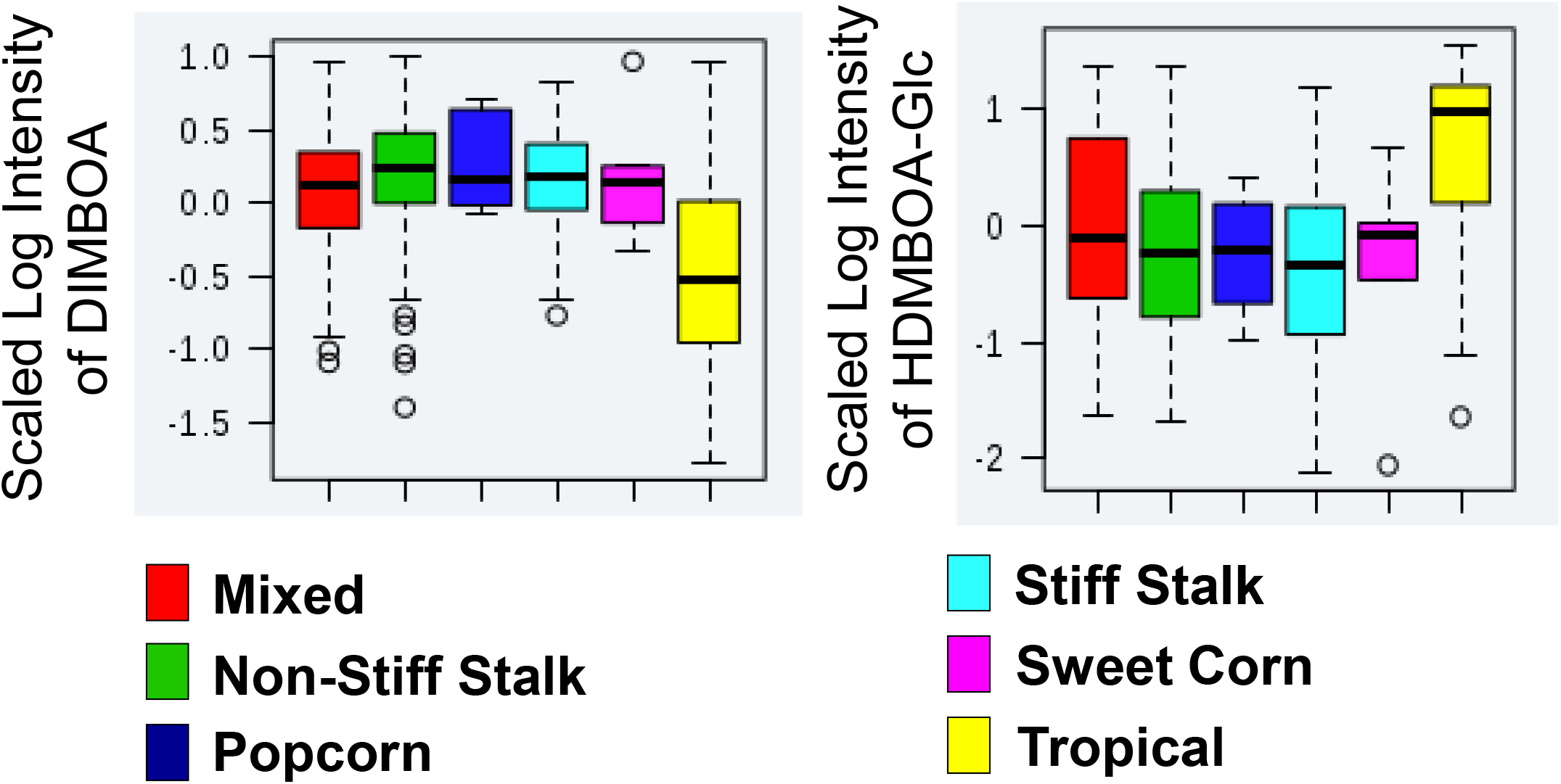
Tropical and temperate maize lines accumulate different benzoxazinoid compounds. Maize inbred lines are assigned to genetic subpopulations defined in Flint-Garcia et al., 2005. The median of each group is denoted by the central think black line, quartiles by the boxes, and 1.5 x the interquartile range by the whiskers.

**Supplemental Figure 5.**
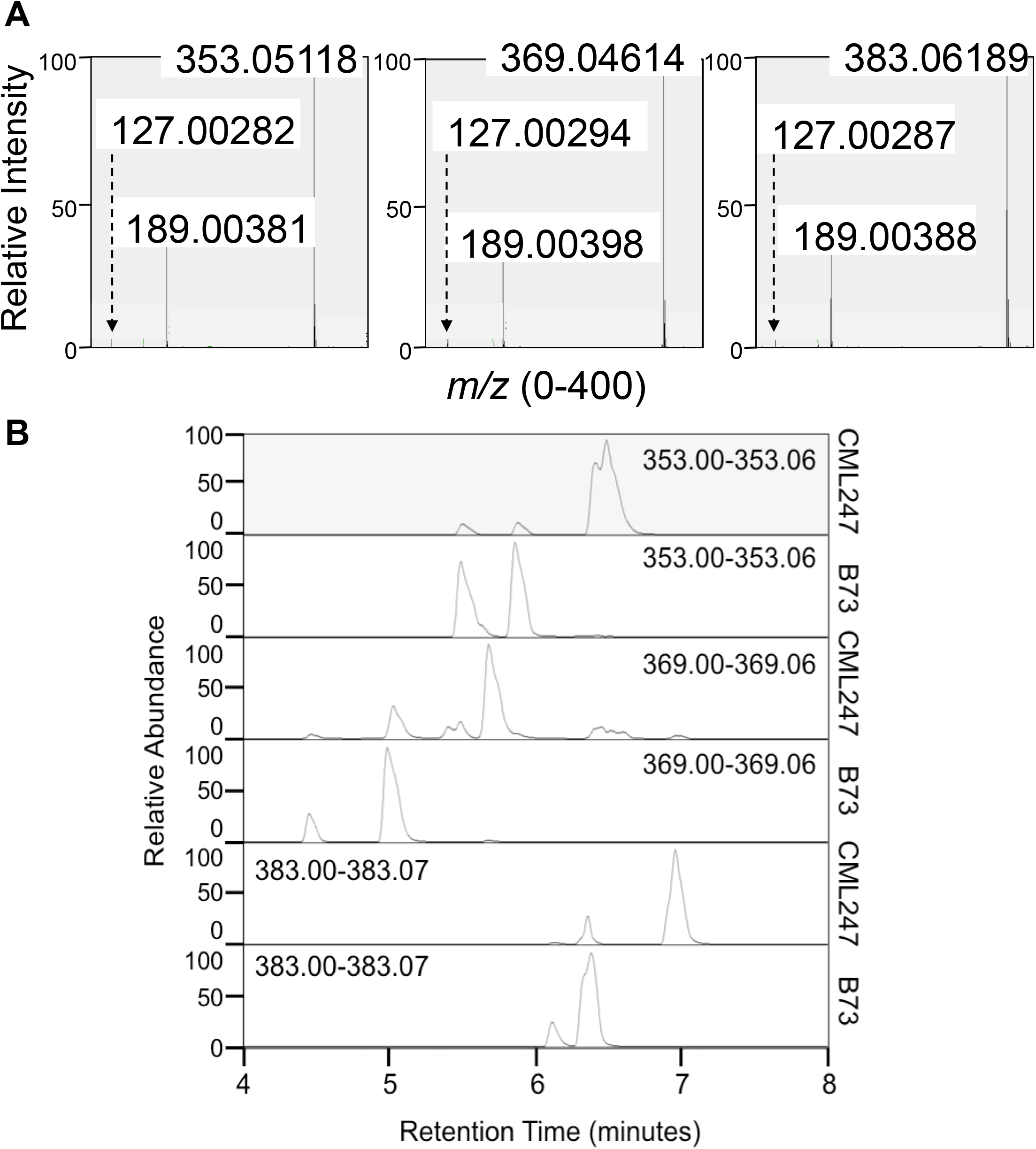
Phenylpropanoid-like mass features co-elute with common daughter ions. (A) Mass spectrum scans of three mass features co-eluting with phenylpropanoid-like UV absorbance peaks are shown. The parental ions and the two shared daughter ions are labeled with their exact *m/z* measurement. (B) In two different maize inbred lines, the predominant peaks at each specific *m/z* range eluting at different retention times likely represent different structural isomers of the same compound.

**Supplemental Figure 6.**
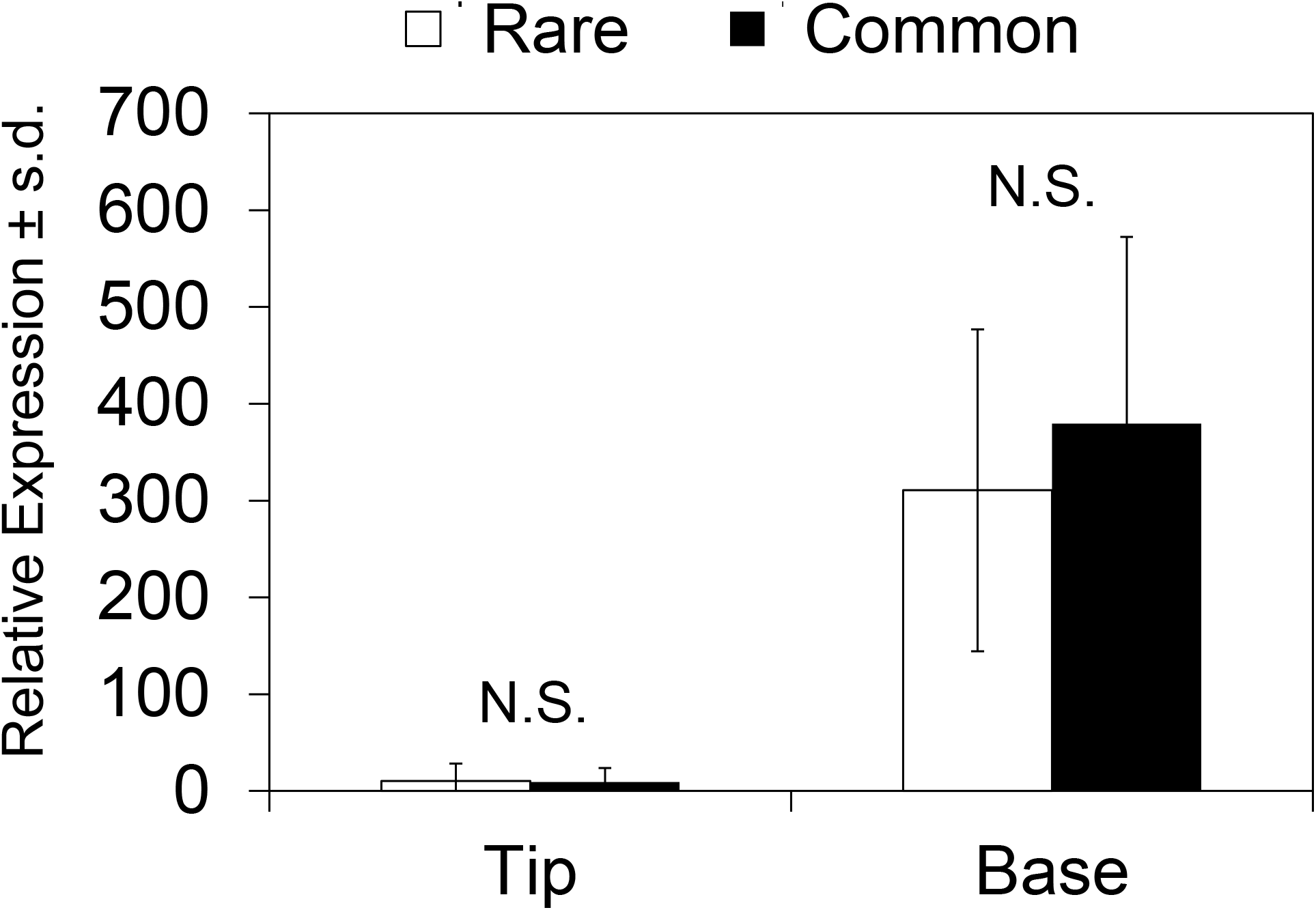
Expression of the citrate synthase family gene is not significantly different between maize inbred lines accumulating different phenylpropanoid hydroxycitric acid isomers. GRMZM2G063909 expression in the two tissue types under study was obtained from Kremling et al. (2018) and compared between maize inbred lines accumulating different phenylpropanoid hydroxycitric acid ester isomers. No significant difference (N.S.) in expression is found in either tissue type (p > 0,05; Student’s *t-*test).

**Supplemental Figure 7.**
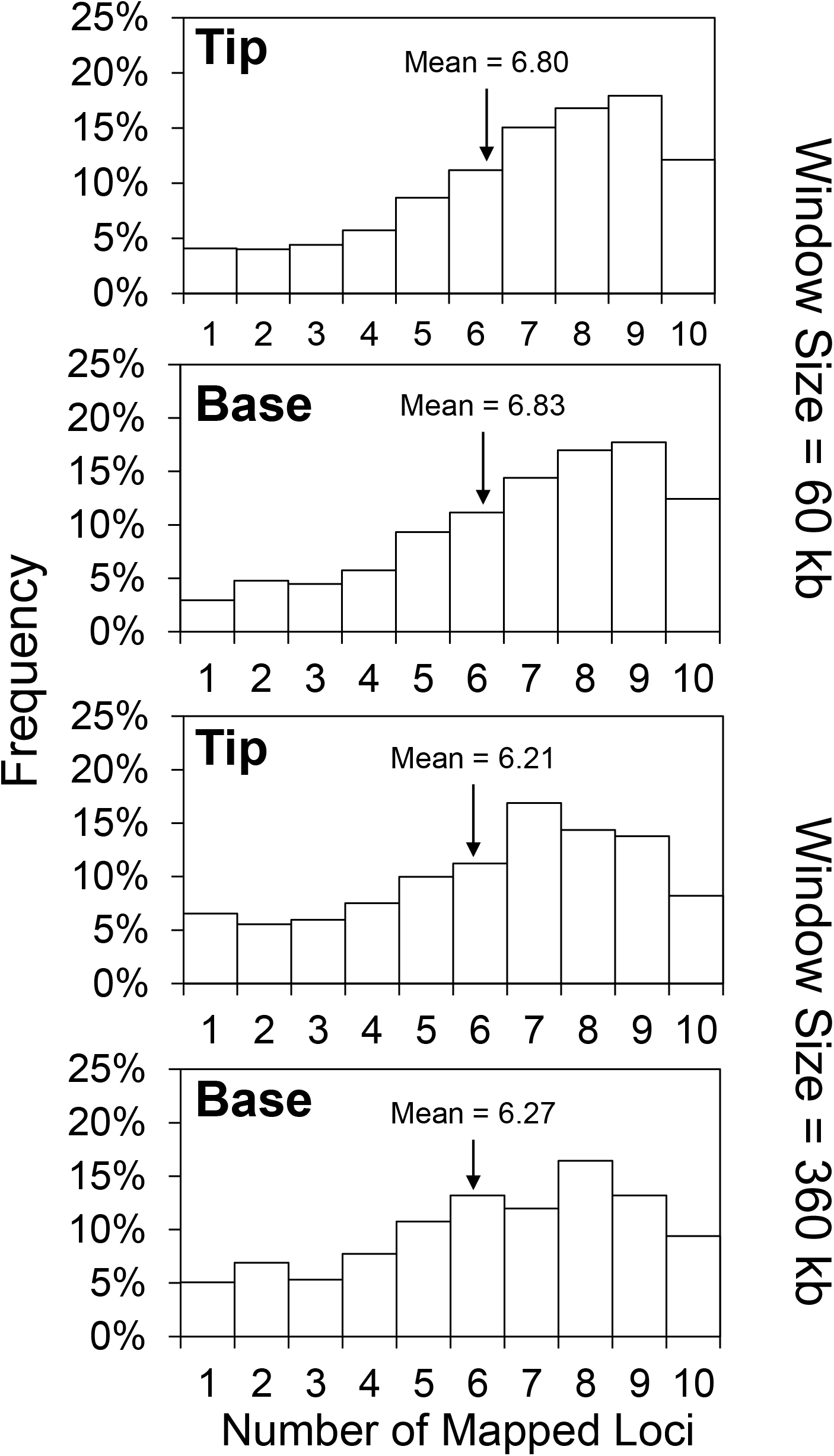
Frequency distributions of genetic architecture complexity of metabolic traits are consistent across different LD window sizes. Distribution of mass features in either tissue type is plotted by number of 60 kbps or 360 kbps LD blocks they are strongly associated with. The statistical mean of each distribution is given and marked by an arrow.

**Supplemental Figure 8.**
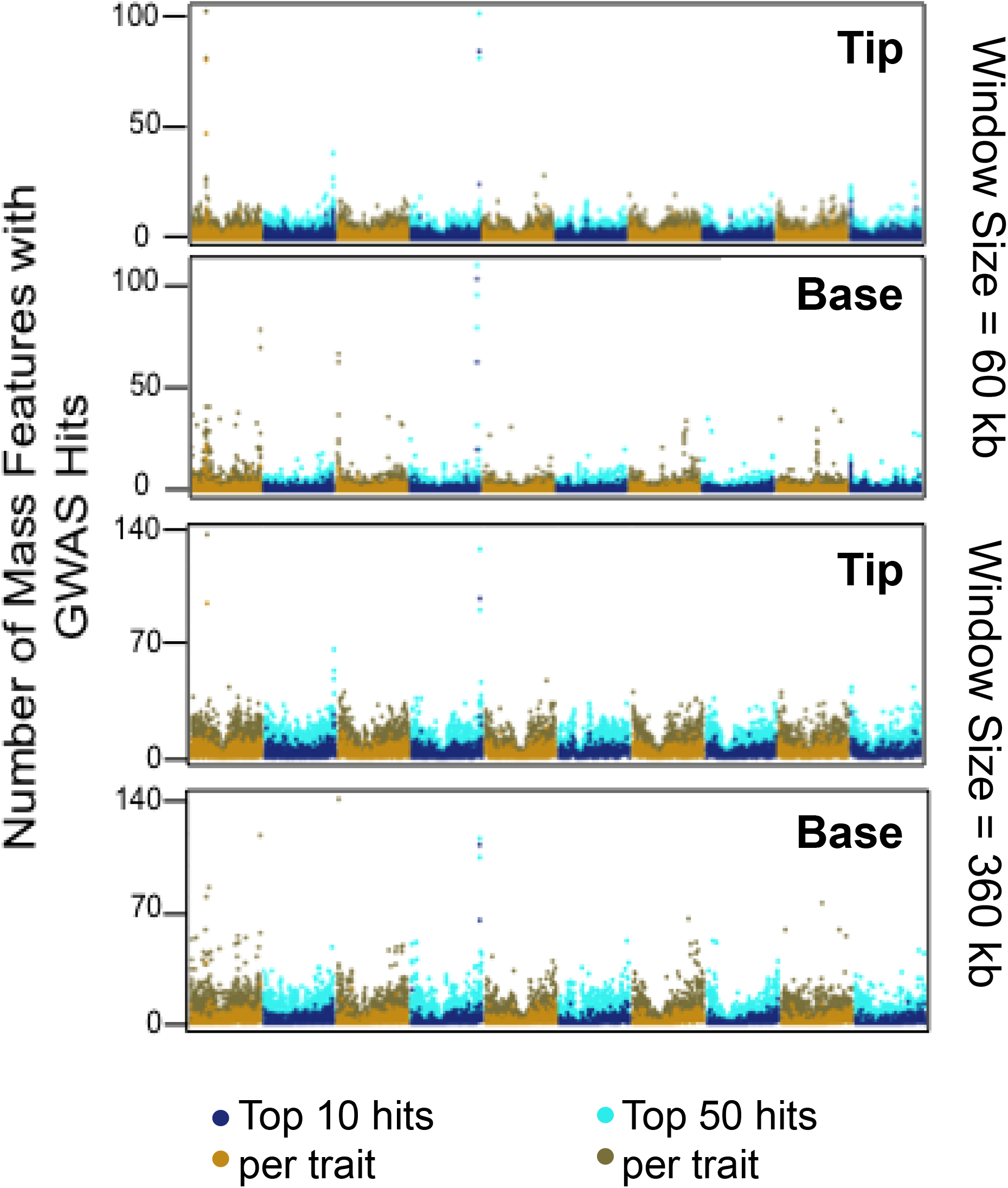
Presence and locations of metabolite QTL hotspots are consistent across different chromosome window sizes. The number of mass features with at least one of their top 10 or top 50 most strongly associated SNP marker located in each 60 kbps or 360 kbps block is plotted for either tissue type.

